# Multiplexing working memory and time: encoding retrospective and prospective information in neural trajectories

**DOI:** 10.1101/2022.07.08.499383

**Authors:** Shanglin Zhou, Michael Seay, Jiannis Taxidis, Peyman Golshani, Dean V. Buonomano

## Abstract

Working memory (WM) and timing are generally considered distinct cognitive functions, but similar neural signatures have been implicated in both. To explore the hypothesis that WM and timing may rely on shared neural mechanisms, we used psychophysical tasks that contained either task-irrelevant timing or WM components. In both cases the task-irrelevant component influenced performance. RNN simulations revealed that cue-specific neural sequences, which multiplexed WM and time, emerged as the dominant regime that captured the behavioral findings. Over the course of training RNN dynamics transitioned from low-dimensional ramps to high-dimensional neural sequences, and depending on task requirements, steady-state or ramping activity was also observed. Analysis of RNN structure revealed that neural sequences relied primarily on inhibitory connections, and could survive the deletion of all excitatory-to-excitatory connections. Our results suggest that in some instances WM is encoded in time-varying neural activity because of the importance of predicting when WM will be used.

## INTRODUCTION

Working memory (WM) refers to the ability to transiently store information, and subsequently use this information in a flexible manner for goal-oriented behaviors and decision making^1, 2^. Timing, here, refers to the ability to track elapsed time after a stimulus, in order to anticipate subsequent external events or generate appropriately timed motor responses^3–5^. While it is widely recognized that the ability to transiently store information about the past and prospectively anticipate external events are among the most fundamental computations the brain performs^1–4, 6–8^, the fields of WM and timing have evolved mostly independently from each other because they have been seen as distinct cognitive functions with different underlying neural mechanisms. Yet, both share critical computational features. Both require transiently storing information, retrospective information in the case of WM and prospective information in the case of timing (e.g., when a delayed reward will occur). In some cases, these properties are mirror images of each other. For example, a timer, such as an hourglass, can be seen as encoding a transient memory that it was recently flipped over *and* of generating a prediction as to when an external event may occur.

Similar neural signatures have been associated with both WM and the encoding of time^3, 5, 9–12^. Although early groundbreaking studies suggested that WM is encoded in steady-state persistent neural activity^13–15^, there is ongoing controversy regarding the neural encoding of WM^9, 10, 16–19^. Broadly speaking, in addition to steady-state persistent activity there are two additional classes of WM models^9, 10, 20^: 1) Time-varying patterns of neural population activity, which can include low-dimensional ramping activity as well as high-dimensional neural trajectories (including, neural sequences); and 2) activity silent mechanisms, in which short-term memory can be stored in the hidden state of neural networks—rather than ongoing spiking activity—through mechanisms such as short-term synaptic plasticity (STSP). Importantly, ramping activity, neural trajectories, and STSP-based changes in the hidden-state of networks have all been proposed to underlie timing as well^3–5, 21^.

The diversity of neural regimes implicated in WM may, in part, be dependent on the presence or absence of implicit timing components. The brain is always attempting to learn the temporal structure of the external world even if it is not explicitly relevant to the task at hand ^7, 22^. Implicit timing enables prediction of when events will take place, thus allowing for preparation and optimal allocation of cognitive resources. Indeed, recent human studies suggest an intimate connection between WM and timing, as WM can be impaired when information has to be retrieved at unexpected times^7, 23^.

We examine the hypothesis that WM and timing are, in some cases, essentially the same computation, and thus rely on the same encoding mechanisms. We first developed two psychophysical tasks that use the same stimulus structure but vary whether the WM or timing components are explicit (required to solve the task) or implicit (task-irrelevant). Participants learned both task-irrelevant WM information during an explicit timing task, and task-irrelevant timing information during an explicit WM task. Given the ongoing challenges in identifying brain regions causally responsible for both the encoding of time and WM, and the success of using artificial neural networks to examine the neural dynamic regimes underlying a diverse set of cortical computations^24^, we trained recurrent neural networks (RNNs) on the same tasks the human participants performed. We show that cue-specific neural sequences emerge as the dominant regime for encoding memoranda and elapsed time from the onset of each memorandum, but overall, that training stages, task structure and hyperparameters captured much of the diversity of the experimentally observed neural dynamic regimes.

## RESULTS

### The differential-Delay-Match-to-Sample and Interval-Stimulus Association tasks

As a first step towards addressing a potential link between WM and timing, we developed variants of the standard Delay-Match-to-Sample (DMS) working memory task. In its simplest form, a DMS task presents either of two cues (in our case, a star or circle denoted by *A* or *B* respectively), and following a delay period either of the two stimuli is presented again, resulting in four conditions (*AA*, *AB*, *BA*, *BB*). Participants are required to differentially respond to the match (*AA*, *BB*) versus nonmatch (*AB*, *BA*) conditions. Typically, the delay between the cue and probe is fixed or randomized, but in our differential-Delay-Match-to-Sample (dDMS) task the cues predicted the delay duration (**Fig. 1a**). For example, the *AA* and *AB* conditions might be associated with a 1 s delay, and *BA* and *BB* trials with a 2.2 s delay—but the delay itself is task-irrelevant. To determine if subjects implicitly learned the temporal structure of the task and whether unexpected delays altered WM performance, the cue-delay contingency was reversed in 20% of the trials (see Methods). The second task (**Fig. 1a, right**), termed an Interval Stimulus Association (ISA) task, was based on the same exact stimulus structure as the dDMS task but framed differently: participants were explicitly instructed to press one key when there was a short delay followed by *A* (Short-*A*) or a long delay followed by *B* (Long-*B*), and another key after Long-*A* or Short-*B* trials. In the ISA task the interval (delay) is explicitly relevant, but the cue (the first stimulus) identity is irrelevant as it just serves as an indicator of t=0 for the interval. Note that during standard trials, the dDMS and ISA tasks are isomorphic—i.e., the correct responses could be produced with either strategy—the difference between the tasks lies in the reverse trials (**Fig. 1a**).

**Fig. 1.**
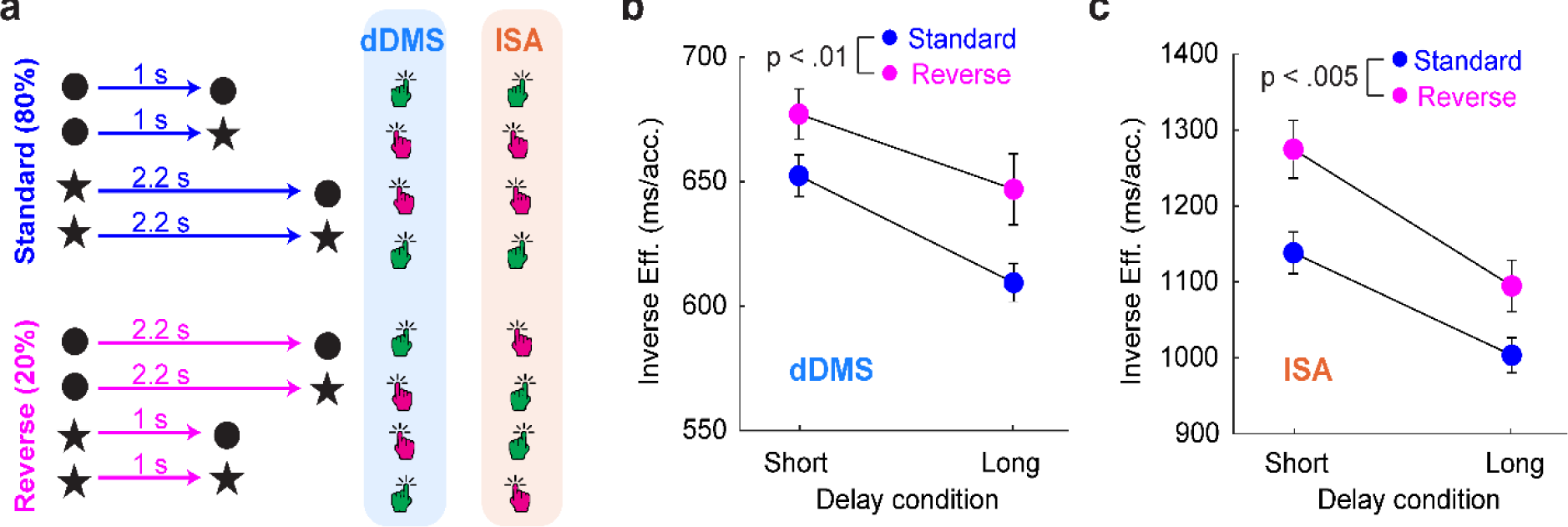
Humans implicitly learn the timing component of a WM task, and the WM component of a timing task. **a,** Schematic of the differential-Delay-Match-to-Sample (dDMS) task, and the explicit timing Interval- Stimulus-Association (ISA) task. Note that the response patterns for the dDMS and ISA tasks only differ during the reverse trials. **b,** Inverse efficiency (RT/accuracy) of human participants on the dDMS task across Standard (cyan) and Reverse (orange) trials. The short and long delays reflect the duration of the actual delay epochs (e.g., a long delay on a Standard trial is an “expected” delay, and a long delay on a Reverse trial corresponds to an “unexpected” delay). There was a significant main effect of Standard vs. Reverse conditions (n = 27, F_1,26_ = 9.05, p<0.01). **c.** Inverse efficiency in the ISA task across Standard and Reverse trials. There was a significant main effect of Standard vs. Reverse conditions (n = 22, F_1,21_ = 11, p<0.005).

To determine if participants implicitly learned the cue-delay associations we analyzed the inverse efficiency (reaction time/accuracy), a measure designed to take into account between-participant differences in speed-accuracy tradeoffs^25, 26^. In the dDMS task (**Fig. 1b**) there was a main effect between Standard-Reverse trials (n = 27, F_1,26_ = 9.05, p<0.01, two-way ANOVA), indicating that the violation of temporal expectation in the reverse trials altered performance. There was also a main effect of the actual delay as expected from the well-known hazard rate effect (F_1,26_ = 11, p<0.001)^27^—after the short interval had elapsed there was increased certainty that the probe will appear at the long delay thus decreasing reaction time (RT). We also examined the raw RT and trial accuracy independently (**Fig. S1**), both exhibited a main effect of Reversal (RT: F1,26 = 7.41, p<0.05; Accuracy: F_1,26_ = 13, p<0.005). To further validate the results of this novel task we performed a replication study (**Fig. S2**), which confirmed a significant Standard-Reverse effect in inverse efficiency (n = 39, F_1,38_ = 8.51, p<0.01), and RT (F_1,38_ = 9.02, p<0.005). There was no main effect of accuracy but there was an interaction between Standard- Reverse and the actual delay (F_1,38_ = 4.1, p<0.05). These results establish that participants implicitly learned the task-irrelevant cue-delay association during a WM task, and that reversing the standard temporal contingency affected WM performance.

We next performed separate experiments using the explicit timing ISA task (**Fig 1c**). Again, there was a significant main effect of Reversal (n = 22, F_1,21_ = 11, p<0.005) on inverse efficiency, as well as on RT (F_1,21_ = 9.2, p<0.01) and accuracy (F_1,21_ = 12.9, p<0.005). A replication study (**Fig. S3**), further confirmed a significant main effect of Standard-Reverse trials on inverse efficiency (n = 25, F_1,24_ = 8.81, p<0.01), RT (F_1,24_ = 9.31, p<0.01), and accuracy (F_1,24_ = 7.78, p<0.05). These results establish that reversing the cue-delay contingency impairs performance on an explicit timing task in which the cue is task-irrelevant, and thus that participants are, in effect, implicitly storing the cue in WM during a timing task.

### Neural sequences in RNNs encoding WM and time

A large body of neurophysiological data across brain areas has revealed a multitude of neural signatures during WM and timing tasks, including neural sequences^12, 28–34^, and firing rate ramps^35–39^. Artificial neural networks, and RNNs in particular, have been invaluable in capturing the experimentally observed dynamics and elucidating the dynamic regimes capable of storing WM and encoding time^11, 40–43^, but to date, with some exceptions^11^, these attempts have primarily focused on either WM *or* timing tasks. Thus, anchored by our dDMS task, we next examined which dynamic regimes emerge in RNNs trained to encode both time and WM (**Fig. 2**). Having established that humans trained on the dDMS task implicitly learn its temporal structure, the RNNs were trained on a timing+WM (T+WM) task, in which the RNN had to learn both the WM and temporal expectation components. RNNs were also trained on two control tasks: a pure WM task without any timing requirements (WM task), and the ISA task which required the RNN to explicitly learn the interval-stimulus association but not the match/nonmatch-to-sample component (ISA task). Note that these tasks do not perfectly parallel the psychophysical studies because the standard distinction between explicit and implicit learning used in the animal literature does not apply to simple RNN models.

**Fig. 2.**
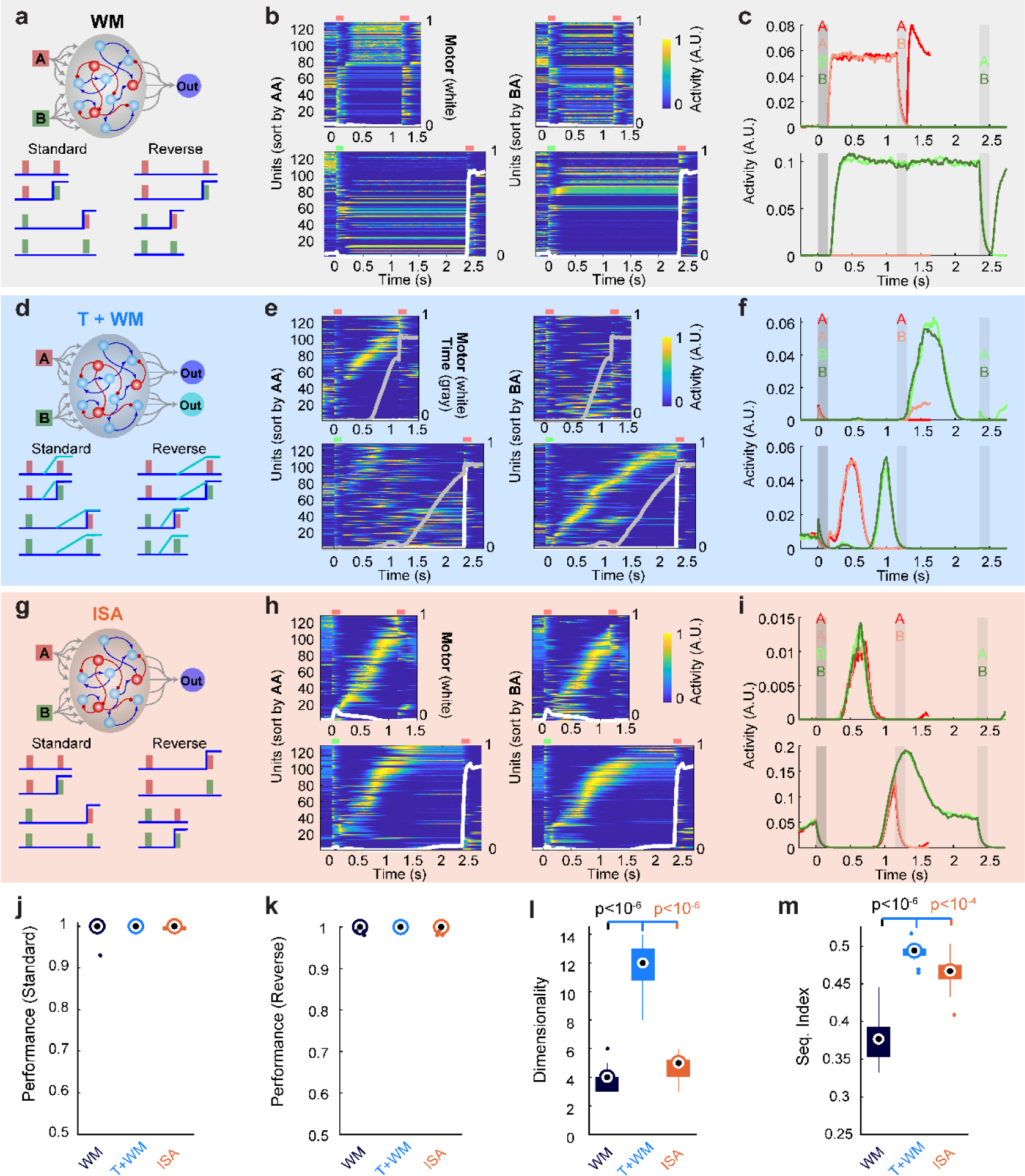
Differential dynamics for the encoding of WM and time across in RNNs trained on three tasks. **a,** Schematic of the RNN architecture and the inputs and target outputs for the WM task during the control and reverse conditions. **b,** Neurograms during the AA (upper row) and BA (lower row) conditions (A=red line above neurogram, B=green), sorted according to the peak time during the short (left) or long (right) delays (standard trials), the images in left and right subpanels of each row is based on the same data, but differentially sorted. The self-sorted neurograms (top left and lower right) are cross-validated (average of even trials sorted on average of odd trials). Only the top 50% of units with the highest peak activity during the delay are shown. Heat map is normalized to one for each unit. The overlaid white line shows the “motor” unit (right y-axis). **c.** Mean activity of two sample units across all four trial types of the standard condition. **d-f,** Similar to **a-c** for the T+WM task. In **e** the activity of the “temporal expectation” output unit is shown in the overlaid gray lines. Note that while each cue elicits a neural sequence, each sequence is different, reflecting the embedding of multiple sequences within the RNN. **g-I,** Same as **a-c** for the ISA task. Note that in this example both cues A and B elicit approximately the same neural sequence (**h**), and the units exhibit similar time fields in response to cues A and B (**i**). **j-m,** Quantification across 17 RNNs trained on the three tasks. Performance (correct match/nonmatch responses) during Standard (**j**) and Reverse (**k**) trials, dimensionality during the delay epoch of the concatenated activity during the short and long trials (**l**), and the sequentiality index during the long trials (**m**). Whisker plots represent medians (circle centers), the interquartile range (boxes), and the most extreme values within 1.5x the interquartile range above or below the interquartile range (whiskers), dots represent “outlier” values.

RNNs were composed of 256 units and had either one (WM, ISA) or two (T+WM) output units. The first output represented the motor response (e.g., nonmatch detection), and the second output represented temporal expectation (implemented as a half-ramp based on anticipatory licking data^31^). The network was composed of three weight matrices: *W^In^*, the input to the RNN; *W^RNN^*, the recurrent weights; and *W^Ou^*^t^, the connections from the RNN to the output. A number of steps were taken to enhance biological realism and improve our ability to dissect the mechanisms underlying the observed network dynamics (see below): 1) Dale’s law was implemented; 2) to capture the low spontaneous activity rates of most cortical neurons a ReLU activation function with a bias of zero was used; 3) in order to focus our mechanistic analyses on the structure of *W^RNN^*, biases of the RNN units and *W^In^* were not trained (see Methods).

The dynamics of the RNNs during the delay period were dramatically different between tasks. RNNs trained on the WM task primarily converged to persistent fixed-point activity during the delay (**Fig. 2a-b**), in which individual units exhibited cue-specific constant levels of activity during the delay epoch (**Fig. 2c**). RNN’s trained on the T+WM exhibited dynamic activity during the delay that when sorted according to latency resembled neural sequences (**Fig 2d-e**). Individual units in these RNNs often exhibited Gaussian-like time fields, in response to one cue and the absence of a response or a different time field in response to the second cue (**Fig. 2f**). The dynamics in the RNNs trained on the ISA task (**Fig. 2g-i**), was more mixed. Specifically, in the example shown in **Fig 2i-h** both cues *A* and *B* triggered the same neural sequence (“erasing” WM information), followed by persistent activity after the initial 1 s period (corresponding to the short delay). This is an effective solution to the ISA task because a categorical encoding of short vs long intervals is sufficient to solve the task.

We compared the performance and dynamics of 17 RNNs trained on the three tasks. Performance, as measured by the correct response of the motor unit, was close to 100% during both the Standard and Reverse trials on all three tasks (**Fig. 2j-k**). To compare the dynamics of the RNNs across tasks we first quantified the effective dimensionality^44^ and the sequentiality index^31^ across the delay periods (see Methods). The dimensionality was significantly higher in the T+WM task compared to both the WM and ISA task (p<10^-6^, Wilcoxon rank sum test), and there was a much smaller difference between the dimensionality for the WM and ISA tasks (p=0.01). The selectivity index was also higher in the T+WM task compared to the WM and ISA tasks (p<10^-6^ and p<10^-4^, respectively), and between the ISA and WM tasks (p<10^-5^). These results are consistent with the interpretation that RNNs converge to fixed-point attractors when they only need to encode WM, but to neural sequences when they need to encode both WM and elapsed time, and to mixed dynamics when they need to encode elapsed time and the nature of the first cue is task-irrelevant (ISA task). We note, however, that there is some variability in the solutions RNNs converged to in each task, particularly during the WM and ISA task. Specifically, sequences could emerge during the WM tasks, while ramping activity and mixed dynamics could emerge in the ISA task (**Fig. S4**).

The T+WM task was designed to capture the human psychophysics data, and critically, in this task two distinct neural sequences generally encoded both WM and timing. Importantly, there is no clear apriori reason that high-dimensional trajectories that approximate neural sequences, should emerge as the dominant solution to encode WM and time. Indeed, intuitively, one might expect much lower dimensional cue-specific ramping activity to encode both WM and time (see below and Discussion).

### Transition from low-dimensional ramping to high-dimensional neural sequences over the course of training

As stated above, either low-dimensional ramping activity or high-dimensional neural sequences can encode both WM and time. Furthermore, both types of dynamics have been observed experimentally during WM or timing tasks^12, 28–33, 35–39^. To date, ramping activity and neural sequences have been treated as fundamentally different dynamic regimes to encode WM or time^3, 9, 11, 45^. To determine if this is indeed the case, we analyzed the development of neural sequence across training in the T+WM task.

Visualization of RNN dynamics from early to late training stages revealed a continuous shift from steady-state activity, to ramps, to neural sequences, both at the population and single-unit level (**Fig. 3a**). Quantified across RNNs this transition was expressed as a progressive increase in the dimensionality of network dynamics (**Fig. 3b**). Importantly, WM performance (i.e., discrimination of match vs. nonmatch trials) peaked very early in training while the dimensionality was fairly low. At these early stages timing as measured by the loss function of the timing output (**Fig. 3c**) or by the ability to decode elapsed time from each cue (**Fig. 3d**), was poor, but increased progressively over the course of training. Finally, there was a strong positive correlation between dimensionality and time decoding performance **(Fig. 3e)**.

**Fig. 3.**
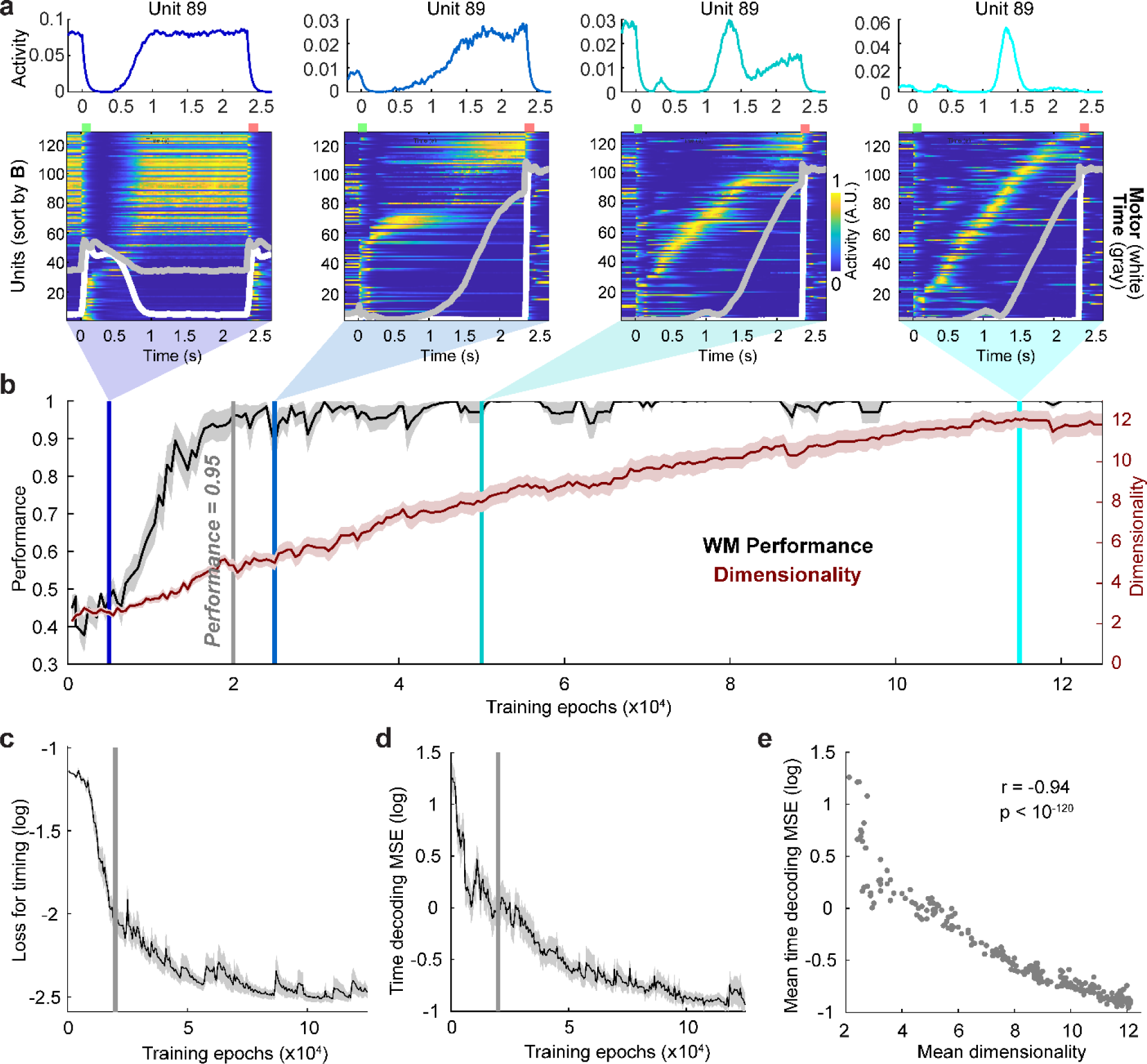
Transition from low-dimensional ramping to high-dimensional neural sequences over the course of training. **a,**Temporal activity profile of a sample unit (top) and neurograms (bottom) of activity in an RNN trained on the T+WM task at the stages of training corresponding to the vertical colored lines in (**b**). Note that in the neurograms the sorting order of the units in the panels is different. **b**, Performance of WM and population dimensionality (during the delay period) across training. The gray vertical line denotes when the mean WM performance reached 0.95. **c**, Learning curve of the loss for the timing output. **d**, Same as (**c**) for MSE of the decoding of elapsed time from each cue. **e**, Relationship between the dimensionality and decoding time MSE averages across 17 RNNs.

The transition from ramps to neural sequences was driven by the timing component of the task, rather than the WM component, because high performance in the match vs. nonmatch discrimination is achieved early in training. These results indicate that from the perspective of an RNN circuit, ramping activity and neural sequences may not represent extremes, but rather a continuum. And additionally, that suggests that in both experiments and computational models dimensionality may be dependent on the degree of training and how well the timing component has been learned.

### Multiplexing of WM and elapsed time

To quantify the ability of the RNNs trained on the three tasks to encode both WM and elapsed time, we used a support vector machine (SVM) to decode both cue and time (Cue-Time)—i.e., elapsed time from the onset of each cue based on population activity. Visual inspection of a sample confusion matrix (predicted versus actual Cue-Time bin) revealed robust Cue-Time decoding in the T+WM task—and thus that either the cue or time could be decoded at any time during the delay (**Fig. 4a**). In contrast, relatively little temporal information was present in the WM task dynamics. And while relatively good decoding of time was possible in the ISA task, ISA-trained RNN’s often confused the cue that signaled the start of each delay. Specifically, the secondary diagonal lines of the confusion matrix indicate that the decoder confused whether time bins of 0.5 – 1 s were associated with Cue *A* or *B*. Across all tasks, the median performance, as measured by the correlation between predicted and actual Cue-Time bins, was above 80% (**Fig. 4b**, left), indicating that even the apparently persistent fixed-point activity in the WM retained a significant amount of temporal information. But decoding was progressively better from the WM, to ISA, to T+WM task as measured both the performance and MSE (**Fig. 4a,** right; all pairwise comparisons were significantly different with p values of at most p<10^-5^).

**Fig. 4.**
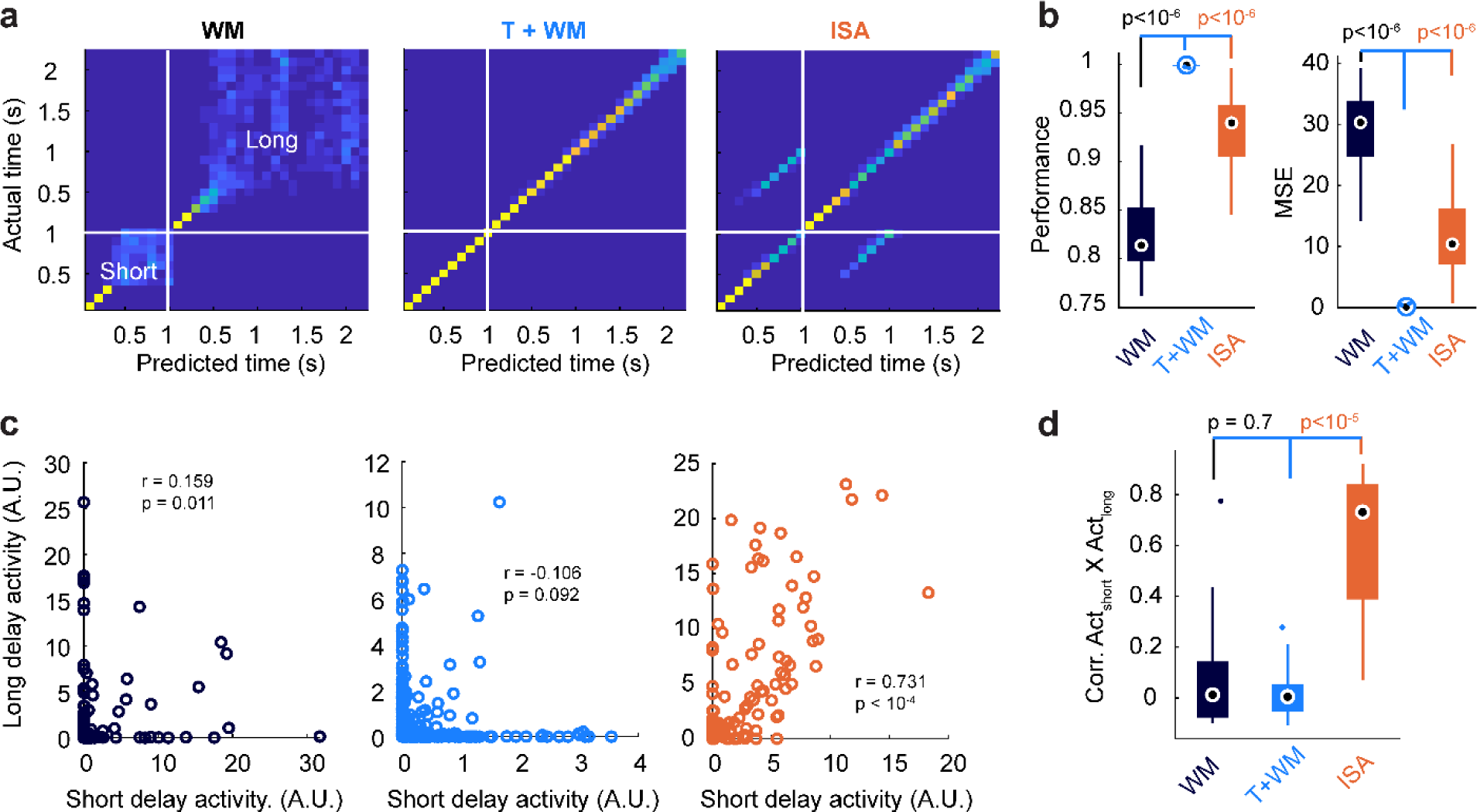
Multiplexing of time and WM. **a,**Confusion matrices of SVM decoding of Cue and time bins (100 ms) for a sample RNN trained on each task. Note that in this example, the ISA network often confuses time bins from approximately 0.5-1 sec during the short (Cue=A) and long (Cue=B) delays and reflected in the off-diagonal bands. **b,** Performance (left) and MSE (right) of the decoders across all RNNs. All pairwise comparisons for both Performance and MSE were significant at p<10^-5^ (Wilcoxon rank sum tests). **c,** Correlation between unit activity for sample RNNs during the short and long delays in the WM (left), T+WM (middle), and ISA (right) tasks. **d.** For the WM and T+WM tasks there was little or no average correlation across all units within each RNN (n=17 in each group). In the ISA task average correlation between unit activity in the short and long delay was high and significantly above the WM and T+WM tasks (p<10^-4^, Wilcoxon rank sum tests).

We next asked if WM memory and time were multiplexed at the level of individual units—as opposed to, for example, a modular strategy in which some units encoded WM and others time. Analysis of the correlation of the mean activity during the delay epochs revealed largely nonoverlapping populations of units activated in response to the short and long cues in the WM and T+WM tasks but largely overlapping in the ISA task (**Fig. 4c-d**). The median Pearson correlation was close to zero across RNNs for the WM and T+WM tasks, but above 0.75 in the ISA task. These results reflect the preservation of cue-specific information during the delay period in the WM and T+WM tasks, but significant loss of cue-specific information during an explicit timing task (ISA). The multiplexing of WM and timing information in the T+WM task at the level of individual units, was confirmed in the high levels of mutual information individual units contained about both cue and elapsed time, as well as the high degree of correlation between them (**Fig. S5**).

### Neural sequences rely heavily on tuned inhibitory connectivity

While dynamic regimes that approximate neural sequences have been observed in many brain areas^12, 28– 34, 46^ and artificial neural networks^44, 47–49^, the circuit mechanisms underlying the production of neural sequences is not understood. This is particularly true in cases that cannot be accounted for by simple feedforward architectures, such as when the same circuit can produce multiple sequences or the units have time fields that rely on circuit rather than intrinsic mechanisms. To dissect the circuit principles underlying the emergence of neural sequences we partitioned *W^RNN^* into its four sub-matrices 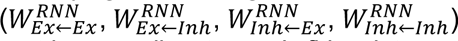, and ordered both the pre- and postsynaptic neurons of the matrix according to peak firing latency during the delay (the long delay was used for visualization purposes). The sorting was performed separately for the Ex and Inh populations (**Fig. 5a**). Next, to extract any general structure underlying the neural sequences we merged the segmented and sorted weight matrices across all 17 RNNs into a master weight matrix (**Fig. 5b-d**). Note that the weights of the entire network are shown, including the units that never fired during the long delay or whose peak was outside the delay, which were placed first in the sorting order—thus it is the weight structure of the latter units of the sorted sequence that are associated with the neural dynamics during the delay epoch. Additionally, the number of units participating in the delay dynamics varied considerably across RNNs. Despite these significant sources of variability, a dominant diagonal component is visible in the synaptic structure of all four weight submatrices in the T+WM task. Critically, for T+WM task there is not a single salient diagonal, and when sorted according to the short-delay another diagonal is visible (data not shown).

**Fig. 5.**
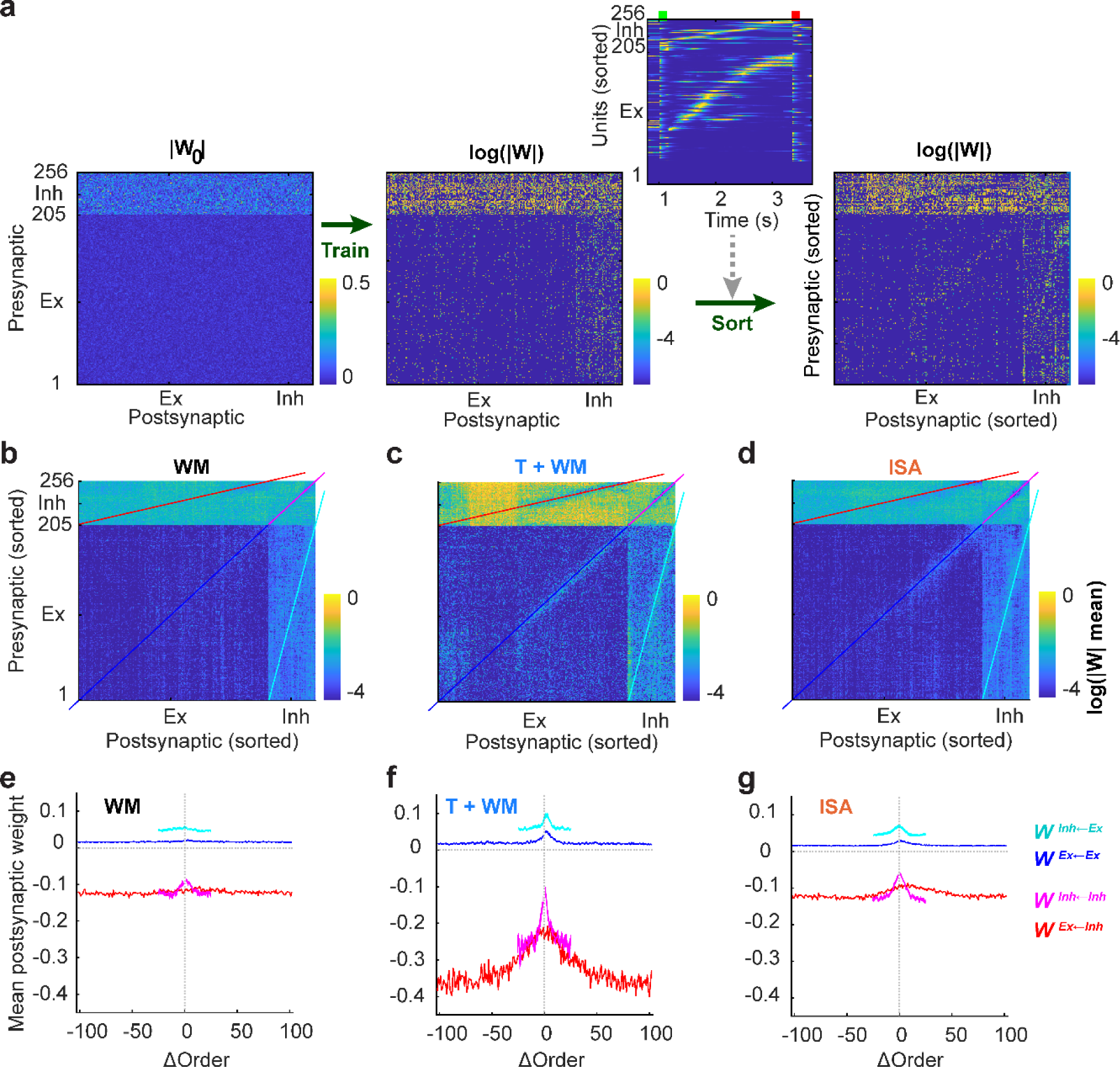
Circuit motifs underlying the generation of multiple neural sequences. **a,** Absolute weights of a sample recurrent weight matrix before training (left), log of weights after training on the T+WM task (middle), and the same posttraining weight matrix sorted by the peak latency times of the excitatory and inhibitory units (inset) during the delay epoch (right). **b-d,** Sorted weight matrices collapsed across all 17 RNNs trained on the WM (**b**), T+WM (**c**), and ISA (**d**) tasks. All matrices are plotted on the same log scale. **e-g,** Mean weights from presynaptic units across all RNNs trained on each task. The presynaptic units are arranged according to their relative peak firing latency during the delay period. Thus ΔOrder=0 captures the mean synaptic weights between pre and postsynaptic units that have the same peak latency, ΔOrder values of 10 and -10 capture the mean weights of the connections from presynaptic units to the postsynaptic units whose peak activity is 10 ms after or before, respectively, the peak of their corresponding presynaptic units.

To quantify the structure of weight submatrices, we averaged the weights according to the relative differences in peak activity latency of the presynaptic units (essentially the average of the main and secondary diagonals in **Fig. 5b-d**), allowing for the visualization of the net synaptic relationships between a presynaptic unit and the postsynaptic units that fired before or after it (**Fig. 5e-g**). In the T+WM task all four weight submatrices revealed peaks centered at approximately zero (corresponding to the main diagonal of the sub-matrices in **Fig. 5c**). The 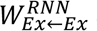 submatrix reveals that presynaptic excitatory units provide above average input to excitatory units that fire shortly before and after it. Critically, the connectivity is asymmetric: stronger in the forward than backward direction. The stronger feedforward influence drives the sequence forward, and the reverse projections contribute to the width of the time field. There were differences in the 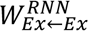 submatrix between the T+WM task compared to the other tasks, but it was the inhibitory connections that were most distinct. For example, the 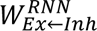 tuning was the broadest and much stronger in the T+WM task. Thus, there was a marked window of disinhibition from inhibitory to excitatory neurons with similar time fields (**Fig. 5f**). To establish a causal link between the structure of each weight submatrix and the observed neural dynamics we examined RNN performance after shuffling all nonzero weights in each submatrix (**Fig. S6**). Shuffling either of the excitatory or inhibitory matrices resulted in catastrophic drops in performance in the WM and ISA tasks. In contrast for the T+WM task shuffling the excitatory weights (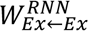 or 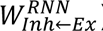) led to median performance levels of approximately 75%, while shuffling the inhibitory weights (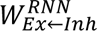 or 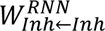) resulted in near-chance performance.

These results indicate that in order for the same RNN to generate multiple neural sequences, it relies more on the connectivity structure of inhibitory connections, than excitatory connections. Specifically, the generation of multiple sequences relies on a transient window of disinhibition, which allows the weak excitation to flow through while globally suppressing the patterns associated with other sequences. These results generate the prediction that the most important site of plasticity for the generation of neural sequences is inhibitory plasticity onto excitatory neurons rather than between excitatory neurons.

### Importance of hyperparameters

To determine if the emergence of neural sequences was dependent on RNN hyperparameters we contrasted the RNN dynamics trained on all three tasks across different hyperparameter configurations (see Methods), including learning rate, L2 activity regularization, noise in the recurrent units, *W^RNN^* initialization, presence or absence of Dale’s law, activation function, and the profile of the temporal expectation function (half- versus full-ramp). Each hyperparameter took on one of two values anchored around the default set of parameters used above, for a total of 288 RNNs over the three tasks and twelve seeds. Across all hyperparameters (**Fig. S7**), with one exception, the dimensionality of the dynamics during the delay period was significantly higher in the T+WM task compared to the WM task (p values of at most 10^-4^, Wilcoxon rank sum test) and the ISA task (p values of at most 0.001). The exception was the use of the softplus activation function versus the default ReLU function. For the softplus function the dimensionality was uniformly low at a value of 2 for all tasks. These data suggest that the use of a softplus function shifts the encoding of time and WM in the T+WM task to ramps rather than population clocks (**Fig. S8**). Indeed, by fitting the activity of the RNN units to both linear ramps and Gaussians during the delay, we observed a dramatic shift in the goodness of fit: while the units from RNNs trained with the ReLU activation function were on average very poorly fit by linear ramps, softplus units were fit quite well (**Fig. S9**). Consistent with the results above, indicating that low-dimensional ramps are not well-suited for flexible timing, the ability of softplus RNNs to generate the half-ramp timing output was worse (the MSE of the timing units was significantly higher compared to the ReLU RNNs; p<10^-4^, Wilcoxon rank sum test). The ReLU activation function is often considered to be highly efficient^50^, and perhaps more naturalistic in that they would seem to better capture the discrete nature of neuronal thresholds and can have output values of zero. The shift from high-dimensional to low-dimensional regimes, may be a result of an interaction between the learning algorithm and the continuous derivative of the softplus function.

As mentioned above, depending on brain area, both the high-dimensional neural sequences and low-dimensional ramps are indeed observed in timing and WM tasks (see Discussion). Thus our results establish that RNNs can account for both these experimentally observed regimes in a hyperparameter- dependent fashion. Raising the possibility that differential intrinsic neuronal properties in different areas could contribute their observed dynamics.

Across all other hyperparameters, the dimensionality and formation of neural sequences in the T+WM task were robust, but we observed that some hyperparameters had surprising effects on the relative contribution of excitatory and inhibitory units. As described above, in the T+WM task the weights from the excitatory units contributed less to the dynamics than that of the inhibitory weights (**Fig. 5** and **Fig. S6**), this phenomenon was further amplified when the noise of the recurrent units (*σ^RNN^*) was increased from 0.005 to 0.05 (**Fig. 6**). At the default noise level of *σ^RNN^* = 0.005, deletion of all excitatory-to-excitatory weights (i.e., zeroing the entire 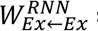 submatrix after training), resulted in a catastrophic drop in performance. Note that even in the absence of excitatory-to-excitatory connections (and biases of zero) some activity is driven by the noise. Surprisingly, at noise levels of *σ^RNN^* = 0.05, ablation of all excitatory-to-excitatory connections, only had a modest effect on the dynamics and performance (median > 90%) during the T+WM task (**Fig. 6c-d**). Critically, in the ISA task deletion of excitatory-to-excitatory weights decreased performance to chance. In other words, there is a fundamental shift in circuit architecture when RNNs encode a single sequence (ISA) versus two sequences (T+WM). In the former case, RNNs rely heavily on excitatory-to-excitatory connections, but in the latter case RNNs rely primarily on inhibitory connections.

**Fig. 6.**
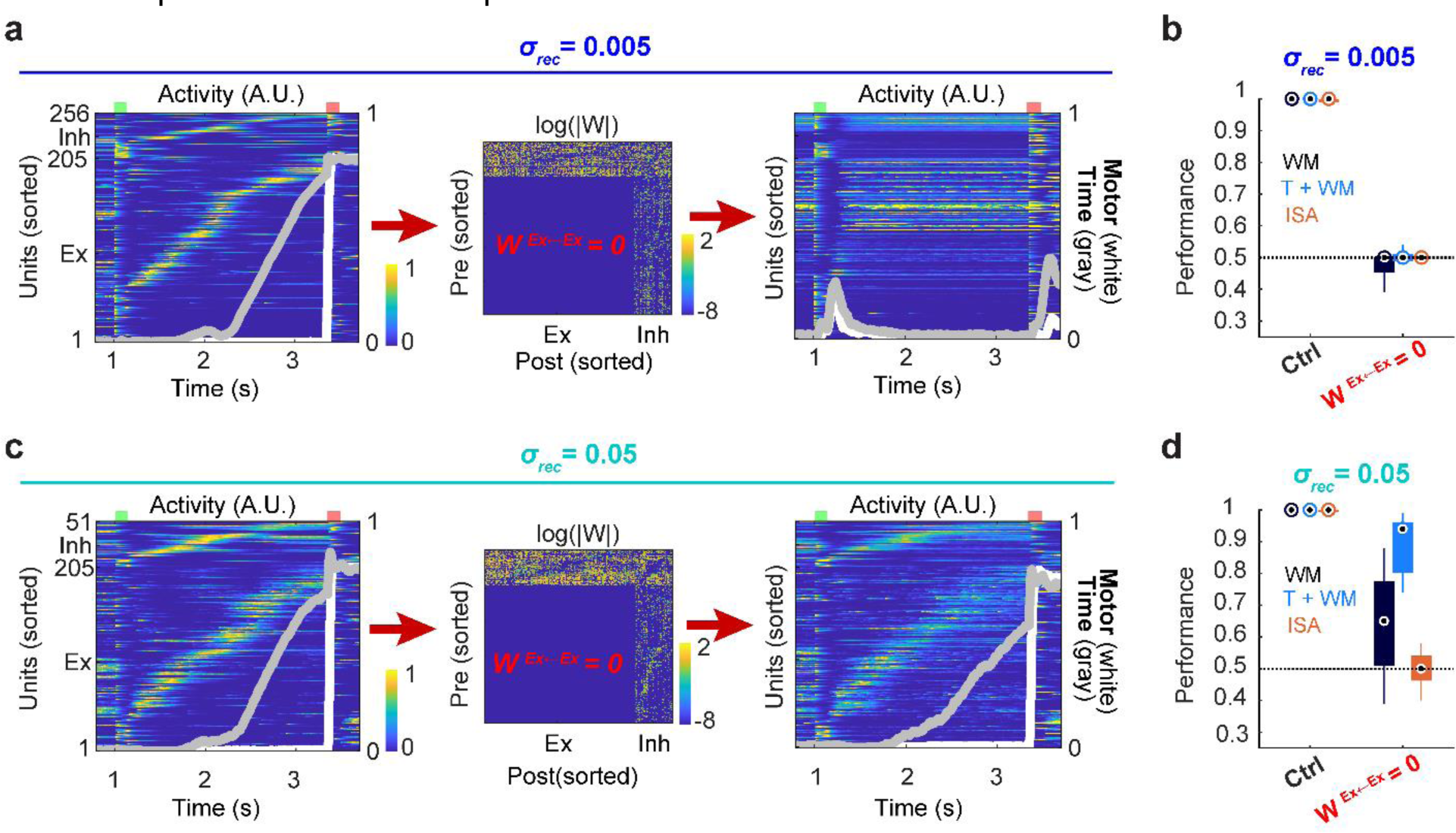
Ablation of all excitatory→excitatory connections has little effect on performance in the T+WM task. **a,**Sample dynamics during the long (BA) trials with default low noise in the RNNs σ=0.005 (left). Dynamics after ablation of all Ex→Ex connections (right). **b,** The mean performance for all three tasks fell from near-perfect to chance upon deleting all Ex→Ex connections. **c,** Dynamics corresponding to panel A in a sample RNN with noise σ=0.05 before (left) and after (right) ablation of all Ex→Ex connections. **d,** In the presence of higher noise levels, the deletion of all Ex→Ex connections had a modest effect on the performance of the T+WM task, and a moderate effect on the WM task. Activity scales are the same in the left and right neurograms of panels **a** and **c**.

In the context of neurobiological circuits our interpretation of these findings is that when encoding cue-specific elapsed time, in some contexts, including high noise, RNNs autonomously converge to circuits architectures that resemble the circuit motifs of the striatum, cerebellum, and CA1, i.e., circuits in which there are no excitatory-to-excitatory connections, which are driven by external input and negative— rather than positive—feedback loops^51, 52^.

### Dynamic attractors

While it is widely accepted that the neural dynamics generated by recurrent neural circuits play a fundamental computational role in WM and timing, the dynamics itself has generally been interpreted in the context of standard dynamical system regimes of fixed-point attractors, saddle points, line attractors, and limit cycles. The high-dimensional trajectories and neural sequences observed in the T+WM tasks do not seem to neatly fit into these classes. But as with standard dynamic regimes, a critical question pertains to the stability of the trajectories, i.e. when perturbed, do trajectories further diverge, remain approximately parallel, or converge back to the original trajectory? To address this question, we performed perturbation experiments during the delay period.

We initially contrasted perturbed and unperturbed trajectories in the presence of frozen noise in RNNs trained on the T+WM task. At the level of individual units, the perturbation immediately altered activity levels, but over the course of hundreds of milliseconds, not only did activity converge back to the unperturbed levels, but converged in a manner that preserved the original temporal alignment (**Fig. 7a**). The effect of the perturbation at the population level can be visualized in the cross-Euclidean distance matrix (**Fig. 7b**), which shows that after the perturbation the main diagonal seems to converge back to close to zero. If the trajectory remained parallel to the original, the diagonal would not return to zero; and if it converged back, but either ahead or behind in time, the minimal values would be off the main diagonal.

**Fig. 7.**
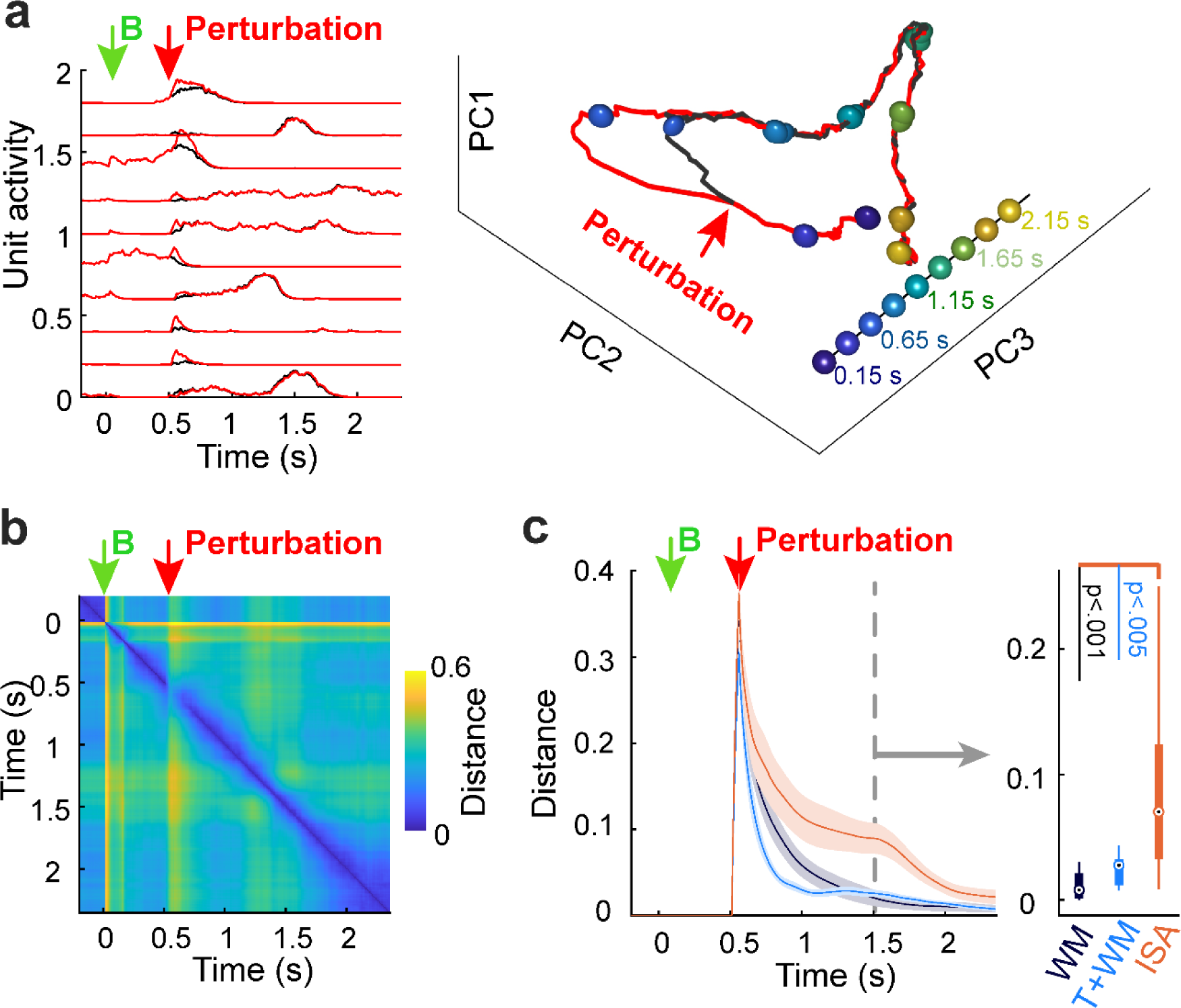
Neural sequences instantiate dynamic attractors. **a,**Activity profile (left) of a sample of 10 units in response to the B stimulus (long cue) in the absence (black) and presence (red) of a perturbation (frozen noise) for the T+WM task, and the unperturbed and perturbed neural trajectories in PCA space (right). Note that the trajectory converges back to the unperturbed trajectory in an approximately time-aligned manner as indicated by the overlapping time marker spheres. **b,** Cross-Euclidean distance matrix between the unperturbed and perturbed trajectories. **c,** Mean Euclidean distance between the unperturbed and perturbed trajectories after the perturbation at 0.5 s averaged across RNNs (left). Median Euclidean distance values (right) 1 s after the perturbation (t=1.5, dashed line).

To quantify the effects of perturbations across RNNs trained on all three tasks we plotted the distance of the main diagonal between the perturbed and unperturbed trajectories (**Fig. 7c**). While the distance does not generally converge to exactly zero, the perturbed trajectory always converges back towards the unperturbed trajectory. These results are consistent with the notion that RNNs trained on the T+WM task implement dynamic attractors, i.e., locally stable transient channels^53, 54^ in which the dynamic attractor is a “hypertube”. Within limits, perturbations of the trajectory can return to the hypertube in motion. Intriguingly, the stability of the neural sequences in the T+WM task was quantitatively comparable to the fixed-point attractor-like dynamics of in the RNNs trained on the WM task. Perturbation analyses based on the direction and magnitude of the vector fields around the trajectories also revealed that the stability of the fixed-point dynamics of the WM task, and the dynamic attractors of the T+WM tasks were comparable (**Fig. S10**).

## DISCUSSION

Working memory and the ability to encode and tell time on the scale of seconds—and thus predict and anticipate external events—are critical to a wide range of cognitive and behavioral tasks. Here we propose that, in some instances, WM and the encoding of elapsed time may be two sides of the same coin, i.e., both WM and elapsed time are represented in the same neural code. This link between WM and time is supported by our findings, and previous findings^7, 23^, that WM is impaired when information has to be recalled at unexpected times, and evidence that in many cases WM is encoded in time-varying patterns of activity^9, 10, 16, 28, 35, 55, 56^.

We first showed that during a WM task (dDMS), participants implicitly learn the task-irrelevant cue-delay associations, and that the time at which the memorandum is accessed not only alters task RT, but accuracy as well. Conversely, during an explicit timing task (ISA), the task-irrelevant cue that marks t=0, also influenced both RT and accuracy. These findings are consistent with results showing that when stimulus-specific temporal structure is present during WM memory tasks, that recall is “dynamically prioritized” ^23^. These psychophysical results, of course, do not establish that WM and time are multiplexed at the neural level, but clearly demonstrate an interaction.

The observation that humans learn the temporal structure of the dDMS task justified training RNNs on a task that required learning of WM and elapsed time (the T+WM task). RNNs are not well suited to study certain psychophysical phenomenon, including implicit learning and behavioral reaction times, as they are not bounded by evolutionarily cognitive strategies or resource optimization constraints. But they have consistently captured the dynamic regimes observed in the brain and provided insights into the biological circuit mechanisms underlying the dynamics^24, 40, 41, 43, 57^. Indeed, consistent with these previous studies, our results revealed RNN dynamics that mirrored a large range of experimental observations.

### Fixed-point and dynamic representations

Early experimental^13, 14^ and computational^58, 59^ studies of WM focused on stable persistent activity, and provided an intuitive computational framework to transiently store memory information in time- *independent* fashion. That is, precisely because information was encoded in a fixed-point attractor, a given memorandum could be retrieved at any time using the same encoding/decoding scheme. A counterintuitive aspect of storing WM in time-varying neural trajectories is that the downstream circuits must recognize that even though the code is changing in time, the memorandum is the same^11^. However, as long as WM is encoded in unique nonoverlapping trajectories, downstream areas can either automatically generalize across time^45^ or learn to recognize the trajectory at all timepoints. Indeed, here, even though the dynamics during the delay period of the T+WM task was time-varying and high-dimensional, performance was near perfect at the learned standard and reverse delays, and generalized to novel intermediary intervals (data not shown).

Dynamic activity has been observed during the delay period in many WM tasks, and it has been shown that the temporal structure of tasks can itself influence the observed dynamics. For example, a task with a fixed delay generated transient dynamics in a premotor area, while random delays resulted in more persistent and stable patterns ^60^. However, in another study the sequentiality of population activity in the PFC was larger in a WM task with a random compared to a fixed delay^61^. One advantage of the dDMS task—in contrast standard DMS tasks in which all stimuli share the same delay—is that if WM is encoded in persistent stable activity, both memoranda should elicit fixed-point dynamics. However, if WM is multiplexed with time, the duration or speed of the dynamics should be distinct across different memoranda.

### Ramps versus Sequences

There is significant ongoing debate regarding the neural regimes underlying both WM and the coding of elapsed time on the scale of seconds. Critically, however, with the exception of stable persistent activity, the candidate mechanisms for both WM and timing are largely overlapping. Ramping activity, neural sequences, complex neural trajectories, and “activity-silent” models have all been raised as possible mechanisms for both WM and timing. In the context of WM, activity-silent mechanisms have generally focused on short-term synaptic plasticity, which can maintain a memory of previous activity in the absence of ongoing neural activity^9, 20, 62^. Similarly, early computational models and subsequent experimental results suggest that short-term synaptic plasticity underlies some forms of sensory timing, by encoding elapsed time in the so-called “hidden state” of neural circuits ^21, 63, 64^.

Neural sequences and high-dimensional activity have been observed across many brain areas during working memory and timing tasks^12, 28–33, 61, 65–67^. Conversely, low dimensional ramping activity has also been observed across areas and tasks^35–39, 68–70^. Furthermore, even within a single brain area different classes of excitatory neurons may differ in the degree to which they encode time and WM^37^. The diversity and complexity of the experimental findings make it challenging to develop area-specific computational models. But here we have shown that depending on task structure and hyperparameters RNNs converge to fixed-point attractors, ramping firing rate, or neural sequences. Critically, we show that from the perspective of the circuitry generating the dynamics, ramps and neural sequences may not represent fundamentally different regimes since there is a transition from ramps to neural sequences over the course of training.

The current study does not speak to activity-silent models, but provides insights to the potential tradeoffs between encoding information in neural sequences and ramping activity. Specifically, despite their apparent complexity and higher dimensionality, neural trajectories approximating neural sequences emerged in a highly robust manner across ReLU-RNNs trained on the T+WM task. This is consistent with previous RNN models trained on WM or timing tasks, in which neural sequences were observed^43, 71, 72^. Other computational studies have observed the emergence of low dimensional dynamics during WM or timing tasks^11, 73^. We show that task structure, hyperparameter choices, and training stages, likely accounts for these differences. A critical question pertains to the computational tradeoffs between the high- and low-dimensional representations of WM and time. One clear tradeoff pertains to ease of generalization to novel delays, and the use of the neural representations by downstream areas to create arbitrary time-varying outputs. While ramping activity in RNNs is a highly limited representation in terms of its ability to generate outputs other than ramps (even including the half-ramp used here), ramps are well suited to temporal generalization^45^. In contrast, neural sequences provide a robust high-dimensional set of basis functions that can drive arbitrarily complex temporal outputs including the default half ramp used here^31^, but require more training in order to generalize to novel time points.

### Conclusions and predictions

Internally generated high-dimensional neural trajectories, including neural sequences, have been reported in a large number of brain areas across many tasks^12, 28–33, 46, 61, 65–67^, and present in many computational models ^43, 44, 53, 71, 74^. We postulate that this is because neural sequences represent a canonical dynamic regime to encode WM, time, and generate motor patterns. The relatively high dimensionality, stability, and quasi-orthogonality of neural sequences, make them well suited for downstream areas to generate either simple or complex time-varying output patterns^31^. Our results predict that neural sequences observed in vivo are not the product of feed-forward architectures as proposed in some models^48, 75^, but require recurrent connectivity. Specifically, sequence generation relies on opening a transient window of disinhibition and allowing non-specific excitation or tonic input to drive the sequences forward^52^. Regarding the biological mechanisms underlying the emergence of neural sequences our results also predict that neural sequences are strongly dependent on inhibitory plasticity, and somewhat paradoxically, can be independent of structured excitatory-to-excitatory connections^57^.

Consistent with the notion that memory serves both retrospective and prospective functions^76–78^, we propose that when WM tasks contain temporal structure, WM and time are multiplexed either in neural sequences or ramping activity. Furthermore, the diversity of experimentally observed neural correlates in WM studies, is in part a reflection of temporal structure of the tasks used. The dDMS task provides a means to address the interaction between WM and implicit timing, as it allows for comparison of the neural dynamics in response to stimuli that have the same WM requirements, but different temporal requirements as to when items in WM will be used.

Multiplexing of WM and time impose additional challenges for downstream decoding^45^, and is unlikely to be a universal encoding scheme for WM. However, multiplexing of WM and time may comprise an effective computational strategy in some instances, because, in addition to the need to transiently store retrospective information the brain is continuously attempting to predict when external events happen, including when WM will be used. Additionally, WM and time may be multiplexed because it provides a learned task-dependent manner to control how long items need to be stored, potentially implementing an expiration time on storage and optimizing cognitive resources.

## Acknowledgements

We thank Ashita Tanwar for assistance with the psychophysics experiments, Jesse Rissman for helpful advice, and Joaquin Fuster for his comments on an earlier version of this manuscript. This research was supported by NIH grant NS116589.

## METHODS

### Human psychophysics

All human psychophysics experiments were approved by the Institutional Review Board of UCLA. Participants provided informed consent before participating and were paid for their participation. Experiments were conducted online, with hosting provided by Gorilla (https://gorilla.sc/) and recruitment provided by Prolific (https://www.prolific.co/). The precision and accuracy of timing on the Gorilla platform (i.e. of visual presentation and reaction time) provides temporal precision with standard deviations of approximately 8-21 ms depending on the exact browser, operating system, and device^79, 80^. Participants accessed the experiment using personal computers running Google Chrome or Mozilla Firefox. No other device types (i.e. phones or tablets) or browsers were allowed. All analyses relied on within subject statistics, thus decreasing the impact of cross-platform variability. Participants on the Prolific platform were only eligible for the study if they were between the ages of 18 and 40, residing in the United States, fluent in English, and had never participated in an online study from our laboratory on Prolific. Before beginning the task, participants read and signed an informed consent form that asked them to: 1) complete the study in a quiet place without distractions, 2) maximize their browser window and not adjust it during the experiment, 3) have normal or corrected-to-normal vision, and 4) not participate if they had a history of seizures, epilepsy, or stroke. After providing consent, participants completed a short demographics form including their age, handedness, and gender. Participants were then given instructions on how to perform the task, which stressed the importance of both speed and accuracy. Participants were also informed that if they were faster and more accurate than the average of the other participants in a given sample of participants, they would receive a bonus payment. Across all experiments 130 participants (62 female, 5 left-handed, mean age = 29, range 18-40) participated in the study. Data from seventeen participants were excluded from analysis due to low accuracy (less than 70%) or consistently slow reaction times (RT) such that too few trials met inclusion criteria (less than 50% of total possible trials in any Reversal x Delay condition remaining after RT exclusion, see below).

#### differential-Delayed-Match-to-Sample (dDMS) task

The background was always white, and all stimuli were black and presented in the center of the screen. First, a 150 ms duration fixation cross was presented, which indicated the start of a new trial. Following a 500-1000 ms interval, a 150 ms duration visual cue was presented, which could either be a black circle or black star, matched for area, with 50% probability. After the delay epoch (see below), a 150 ms duration probe stimulus was presented that was either the same or remaining stimulus with 50% probability. Participants were instructed to press one of two buttons on their keyboards, F or J, to indicate whether they thought the cue and probe stimuli matched or did not match (counterbalanced across participants). The response period was unlimited in duration, and the task did not proceed unless a response was given. All incorrect responses were followed by negative feedback (a “thumbs down” icon). After each response there was a 1500-2000 ms intertrial interval.

The critical manipulation was the delay time, i.e., the interval between the cue (first stimulus) offset and probe (second stimulus) onset. When appearing as a cue, one stimulus (e.g. the circle) was followed by a delay of 1 sec on 80% (“Standard”) of the trials, and a delay of 2.2 sec on the remaining 20% (“Reverse”) of the trials. The other stimulus (e.g. the star) was followed by a 2.2 sec delay on 80% (“Standard”) of the trials and a 1 sec delay on 20% (“Reverse”) of the trials. The mapping between the cue stimulus and the likely memory delay was counterbalanced across participants.

Five blocks of 80 trials (64 Standard, 16 Reverse) were presented for a total of 400 trials. In each block trial order was pseudorandomized with the following constraints: 1) The first eight trials of each block were always standard trials. 2) A Reverse trial could not immediately follow another reverse trial. Participants were given eight standard practice trials with each cue before the first block. Participants were given the opportunity to take short breaks between each block. Each block took approximately eight minutes to complete, and participants finished the experiment in 45 minutes on average. A replication study of the dDMS task (**Fig. S2**) was preregistered (doi.org/10.17605/OSF.IO/XK3JH).

#### Interval-stimulus association (ISA) task

The interval-stimulus association (ISA) task was identical in stimulus structure to the dDMS task, but rather than being instructed to compare the cue and probe stimuli to each other, participants were instructed to make a decision based on the probe stimulus and the delay: e.g., press the F key in response to a short delay followed by a circle or a long delay followed by a star (Short-Circle or Long-Star), and press the J key in response to a long delay followed by a circle or a short-delay followed by a star (Long-Circle or Short-Star). The mapping between the response button and the pair of opposing interval-probe combinations was counterbalanced across participants. Participants were instructed that for any given trial the cue stimulus could be either a Circle or Star and that the cue stimulus was irrelevant to the task beyond indicating the onset of the delay interval. But as in the dDMS task, the cue stimulus identity (Circle or Star) predicted the delay interval on 80% of the trials (Standard trials), while for the remaining 20% of the trials, the cue stimulus was followed by the other delay interval (Reverse trials).

#### Analysis and Statistics

Trials with RTs outside of the range of 100-3000 ms were discarded. Three measures of performance were used: accuracy (percent correct), RT, and the inverse efficiency score (IES). For each condition for every participant, trials with RT values larger than four standard deviations away from the mean were discarded. RTs were calculated as the median of the remaining trials for that condition. The inverse efficiency score (IES), a combined measure of speed and accuracy in which larger values indicate worse performance, was calculated as the median RT divided by accuracy.

Statistics were based on within-subjects 2 x 2 ANOVAs with a Reversal (Standard vs. Reversal trials; e.g., Circle→Short/Star→Long versus Circle→Long/Star→Short) factor and the actual delays (Short vs. Long) factor.

### Recurrent Neural Network Model

#### RNN architecture and training

RNNs were composed of 256 units, an input layer, and an output layer composed of one (WM and ISA tasks) or two (T+WM task) units. The dynamics of the default RNN were described by:

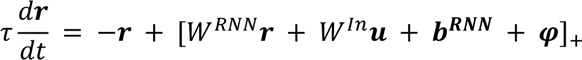

Where ***u*** is the input vector, and ***b****^RNN^* is the bias of the units. Each unit received private Gaussian noise 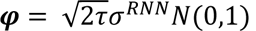,, where *N(0,1)* represents a normal distribution with a mean of 0 and a standard deviation of 1. The threshold linear function [ ]+ represents a ReLU function in which all negative values become 0.

The input layer was composed of 32 inputs representing a range of 0 to 2π. Stimuli *A* and *B* were represented by non-overlapping patterns of activity centered at 1 and 5.2 (corresponding to center activation at units 6 and 28). These patterns can be interpreted as retinotopic or tonotopic activation of two visual or auditory stimuli. The output units (***z***) were nonlinear readouts of the recurrent network:

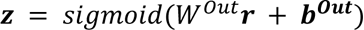

Dale’s law was implemented by assigning 80% (205) of the units as excitatory and 20% (51) as inhibitory. After initializing *W^RNN^*to a random orthogonal matrix (gain = 0.5), the absolute weights of all presynaptic excitatory units were multiplied by 1.0, and those from the inhibitory units were multiplied by 4.0 (to maintain an approximate excitatory/inhibitory balance). During training all weights were clipped at zero during any zero-crossings. The weight matrix was multiplied by a diagonal matrix composed of 1’s and -1’s, corresponding to the excitatory and inhibitory units, respectively^81^.

To enhance our ability to dissect the circuit mechanisms underlying the observed dynamics, the recurrent biases were set to zero, and neither the recurrent biases or the *W^In^* matrix was trained. This approach is consistent with the reverse hierarchy theory ^82^, and the notion that synapses higher in the processing hierarchy are more plastic. Additionally, this approach ensured that the differential RNN dynamics across all networks could be attributed to *W^RNN^*, facilitating the synaptic structure analyses (**Fig. 4**). Thus, only the *W^RNN^*, *W^Out^*, and ***b****^Out^*, parameters were trained.

During training a first-order Euler approximation was used with a *τ* of 50 ms and a *dt* of 10 ms, resulting in a discretization factor *α* = *dt*/*τ* = 0.2. RNNs were trained with ADAM and a batch size of 32. The loss function to be minimized was:

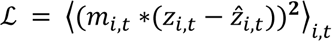

where ℒ represents the time and output unit averaged loss function. *Z*_*i,t*_ and *Ẑ*_*i,t*_ represent the target and actual activity, respectively, of an output unit *i* at time *t*. *m* represents a cost mask that differentially weights the contribution of output units to the loss function during different points in time. The motor output target (*z_1,t_*) was a step function from 0 to 0.8 at probe onset of the nonmatch trials, and when present, the temporal expectation output (*z_2,t_*) was a linear “half” ramp from 0 to 0.8 starting at 50% of the delay period until onset of the probe. For the motor output *m_1,t_* was equal to 2 from 250 ms before onset of the cue stimulus until the onset of the probe stimulus, and 5 during the probe until 500 ms after probe offset (to place a higher weight on the match/nonmatch response), with a grace period 5*dt* (*m_1,t_*=0) during the onset of the probe. For the temporal expectation output, the cost mask was always *m_2,t_* = 1. As with the psychophysics experiments, RNNs were trained on both the Standard and Reverse trials, but reversal trials comprised 10% of the total rather than 20%, to accelerate convergence (note that for the T+WM task the Reverse trials impose a “moving target” for the timing output pattern). Training was stopped when the loss reached 0.0015 for T+WM and 0.001 for the WM and ISA tasks, or reached a total 125,500 update epochs. Different stop criteria were used for T+WM task because the reverse trials make it impossible to fully converge to the same loss values. RNNs were implemented in TensorFlow and based on Yang et al.^41^. Code is available at https://github.com/BuonoLab/Timing-WM_RNN_2022.

During training the onset time of the first stimuli was uniformly varied between 250 and 1000 ms on each trial. The standard delays were 1.0 (short) and 2.2 (long) s. During training these delays were jittered by ±10%, approximately corresponding to psychophysically observed Weber fractions^83^. It is important to note that the presence of “temporal noise” in the form of onset and delay time variability, contributes to the robustness of the solution, and that “spurious” solutions that do not generalize to different onset times or delays can emerge in the absence of this “temporal noise”.

#### Dimensionality

To estimate the dimensionality of the dynamics during the delay periods we first concatenated the average activity of all units during the short and long delays for a final matrix of 256 units x 320 time bins. Concatenation is important to distinguish regimes in which both cues elicit similar sequences (e.g., in the ISA task) versus cases in which both cues elicit distinct sequences (e.g., in the T+WM task). Effective dimensionality was defined as the minimum number of principal components that captured at least 95% of the variance of the activity across time bins^44^.

#### SVM Decoding

For the decoding of Cue and elapsed time, the mean activity of each unit within 100 ms bin was used, comprising a total of 32 (10 and 22 bins for the short and long delays, respectively) input vectors of size 256 per trial. Thus target values represented Bins 1:10 for Cue *A* and Bins 1:22 for Cue *B*). SVMs were trained (SVMTRAIN from LIBSVM 1.2 for Matlab) using multiclass classification, a linear kernel, and a cost parameter of 100. The short and long delay data sets consisted of 25 delay epoch trials of the *AA* and *BA* conditions, and testing relied on leave-one-out cross-validation across the 25 possible replications. Performance was quantified as the correlation between the predicted and target bins, as well as the mean squared error (MSE).

#### Mutual Information

To estimate the per unit mutual information about the cue (i.e., whether the first stimulus was *A* or *B*), the activity was averaged from 100 ms after the end of the cue (to allow for decay of stimulus-evoked activity) to the end of the delay period for each trial. Activity levels were categorically binned into 10 bins from 0 to maximal activity for each unit. Mutual information was calculated across 25 trials of Cues *A* and *B*. To calculate mutual information across time, activity was averaged across ten evenly spaced time bins across the delay period, and again activity was categorically binned into ten activity levels. Maximal mutual information was 1 and 3.32 bits for the Cue and time mutual information estimates, respectively.

#### Perturbation experiments

To test the robustness of RNN dynamics to perturbations, we introduced activity to an input unit during the delay epoch to mimic a distraction produced by an irrelevant stimulus. Specifically, the perturbation input was at π for the input topological position (corresponding to center activation at 17^th^ input) with random weights to the recurrent units similar to the standard inputs. Unless otherwise specified, the perturbation was introduced with the amplitude of 0.25, 500 ms after the onset of the first stimulus for a total duration of 50 ms. Control and perturbed trajectories were obtained using the same noise matrices across units and time (“frozen” noise).

#### Visualization of the velocity fields

To further determine if the dynamics in the T+WM task is consistent with a dynamic attractor regime, we directly quantified the velocity fields around the learned neural trajectories. At each time point *t* on a given neural trajectory (*r(t)*) during the delay period, probe points were sampled from a disc with radius of 0.25 around *r(t)* orthogonal to the tangential direction of the trajectory at time *t*. These probe points were then fed into Equation 1 to obtain the corresponding velocity vectors. Angles between the velocity vectors and tangential/radial direction to the trajectory, along with the amplitude of the vectors were then computed and visualized.

#### Statistics

Comparison across RNN tasks relied on the nonparametric Wilcoxon rank sum test (*ranksum* command in Matlab).

## SUPPLEMENTARY INFORMATION

**Fig. S1.**
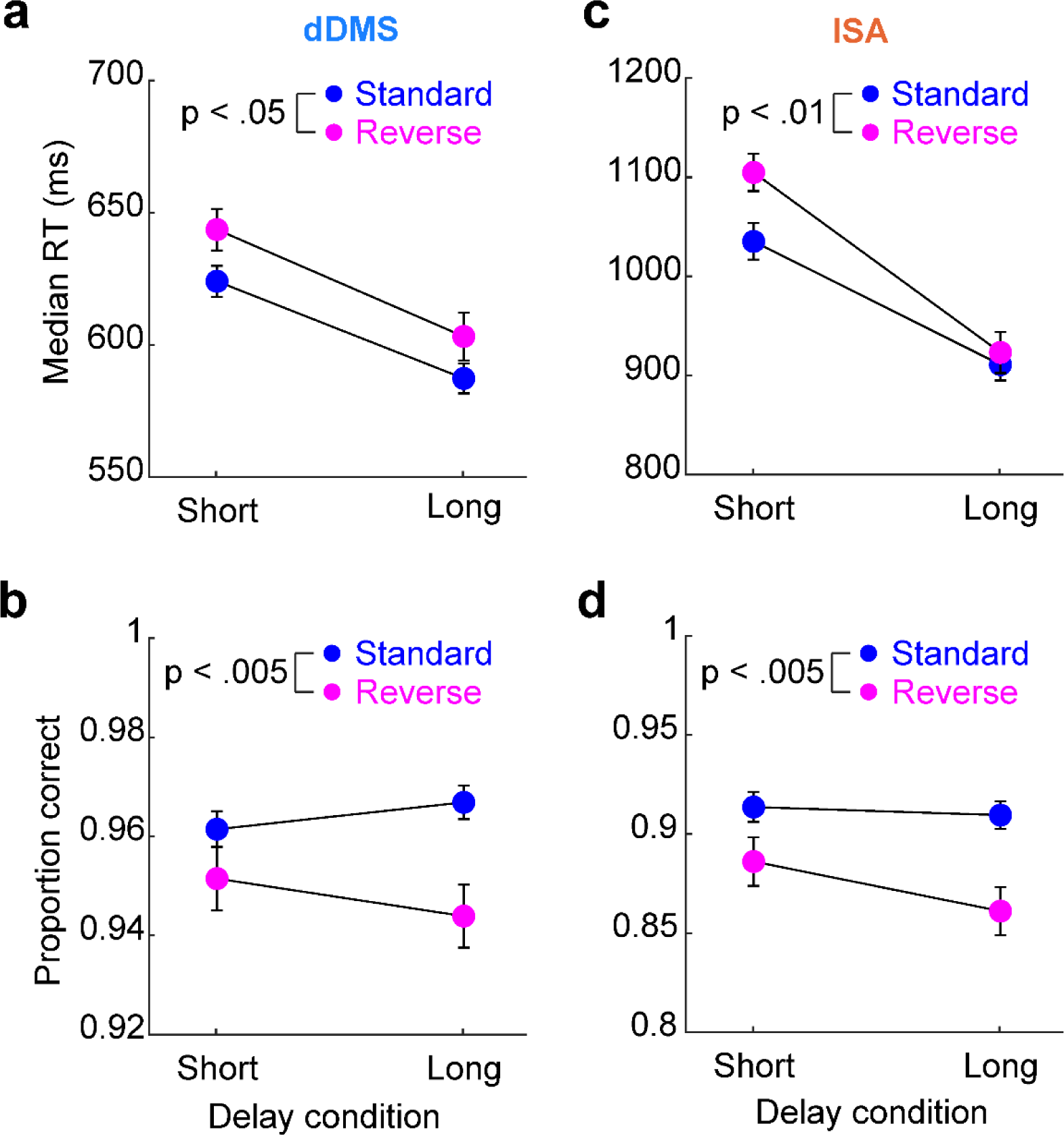
Reaction time and accuracy data for the dDMS and ISA tasks (Fig. 1). **a,** In the dNMS task there was a significant main effect of reversal (Standard vs. Reverse) on reaction time (F_1,26_=7.4, p<0.05; no interaction). **b,** For the dDMS tasks there was also a main effect of reversal on accuracy (F_1,26_=13.4, p<0.005; no interaction). **c,** In the ISA task there was a main effect of reversal (Standard vs. Reverse) on reaction time (F_1,21_=9.2, p<0.01; no interaction). **d,** For the ISA tasks there was also a main effect of reversal on accuracy (F_1,21_=12.9, p<0.005; no interaction). These plots correspond to the same set of experiments shown in Fig. 1b and 1c.

**Fig. S2.**
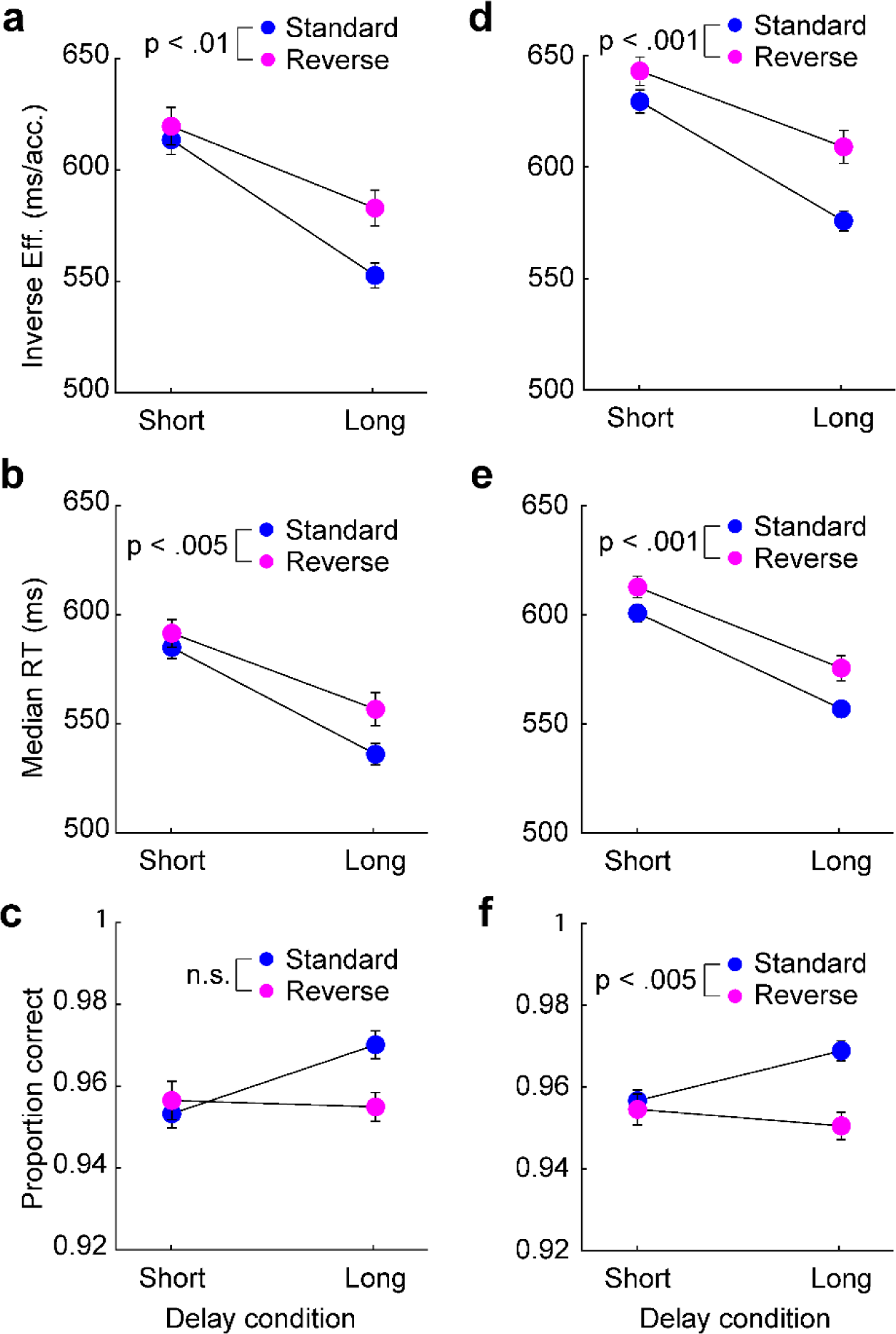
Replication of the dDMS task. **a,** As in the first study (Fig. 1b), an independent replication of the dDMS task revealed that the inverse efficiency score was worse (larger) during the reverse condition (F_1, 38_=8.5, p<0.01; no interaction). There was also a significant main effect of delay (F_1, 38_=36.1, p<10^-5^) reflecting the hazard rate effect. **b,** In the dDMS replication there was also a main effect of reversal on reaction time (F_1, 38_=9.0, p < .005; no interaction). **c,** There was no main effect of reversal on accuracy but there was a significant interaction between the Standard/Reverse conditions and delay (F_1, 38_=4.1, p < .05), again indicating that the implicit learning of the cue-delay associations affected accuracy, but primarily during long delays. **d-f,** A three-way ANOVA that incorporated experiments (original/replication) as a third factor did not reveal a main effect of experiment (or interactions), thus we performed an analyses collapsed across both experiments. There was a significant main effect reversal on inverse-efficiency (F_1, 65_=17.5, p<0.001), reaction time (F_1, 65_=16.6, p<0.001), and accuracy (F_1, 65_=9.0, p < 0.005).

**Fig. S3.**
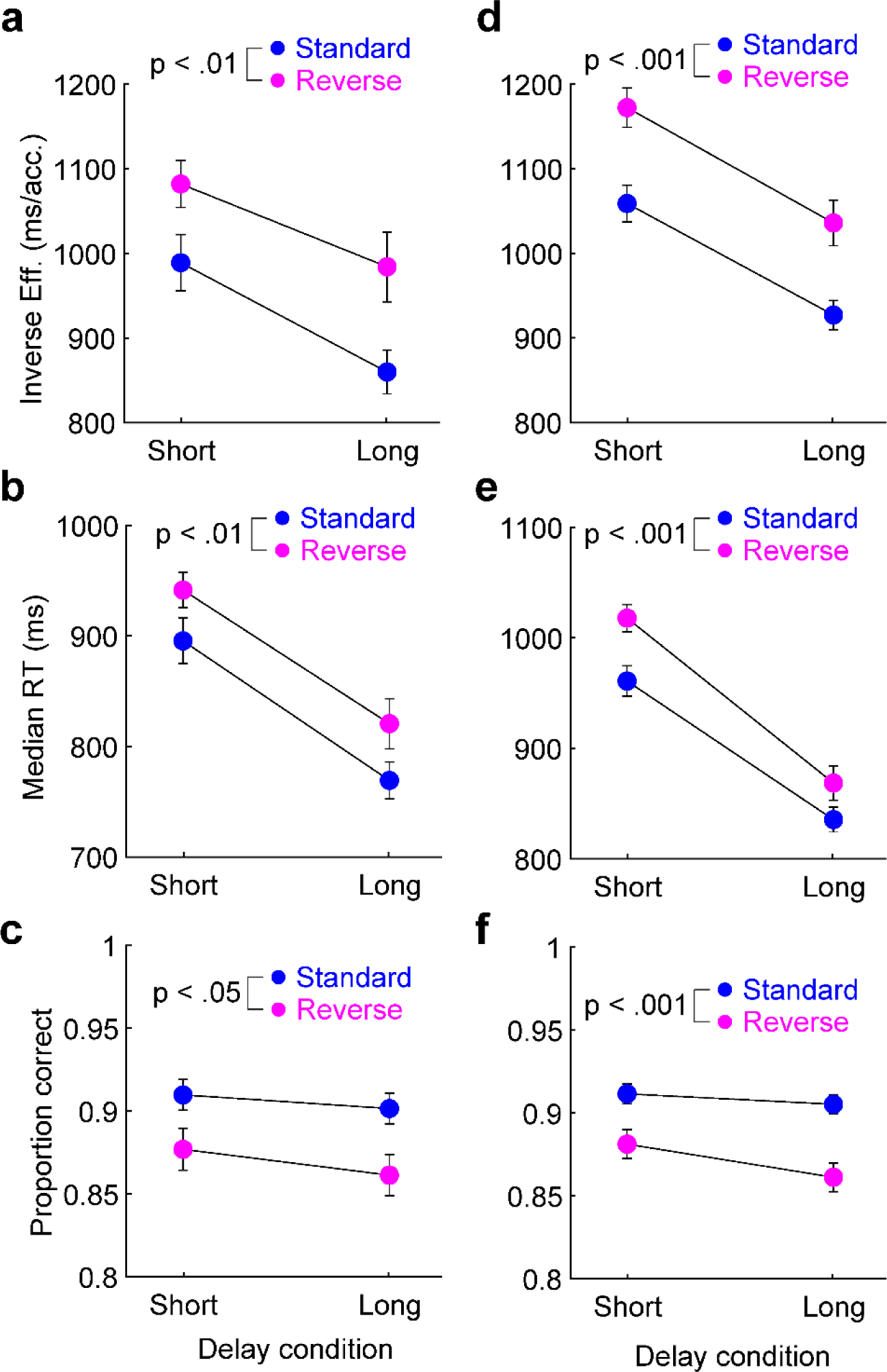
Replication of the ISA task. **a,** Consistent with the first ISA study (Fig. 1c), an independent replication of the ISA task revealed a significant main effect of reversal (Standard vs. Reverse) on inverse- efficiency (F_1,24_=8.8, p <0.01; no interaction), as well as a significant main effect of delay (F_1,24_=5.2, p<0.05) that reflects the hazard effect. **b,** In the ISA replication there was also a main effect of trial type on reaction time (F_1,24_=9.3, p<.01; no interaction). **c,** A main effect of reversal was also present on the accuracy measures (F_1,24_=7.8, p<0.05). **d-f,** A three-way ANOVA that incorporated both ISA experiments (original/replication) as a third factor did not reveal a main effect of experiment (or interactions), thus we performed an analyses collapsed across both experiments. There was a significant main effect of trial type on inverse-efficiency (F_1,46_=19.7, p<.001), median reaction time (F_1,46_=18.5, p<0.001), and accuracy (F_1,46_=19.4, p<0.001).

**Fig. S4.**
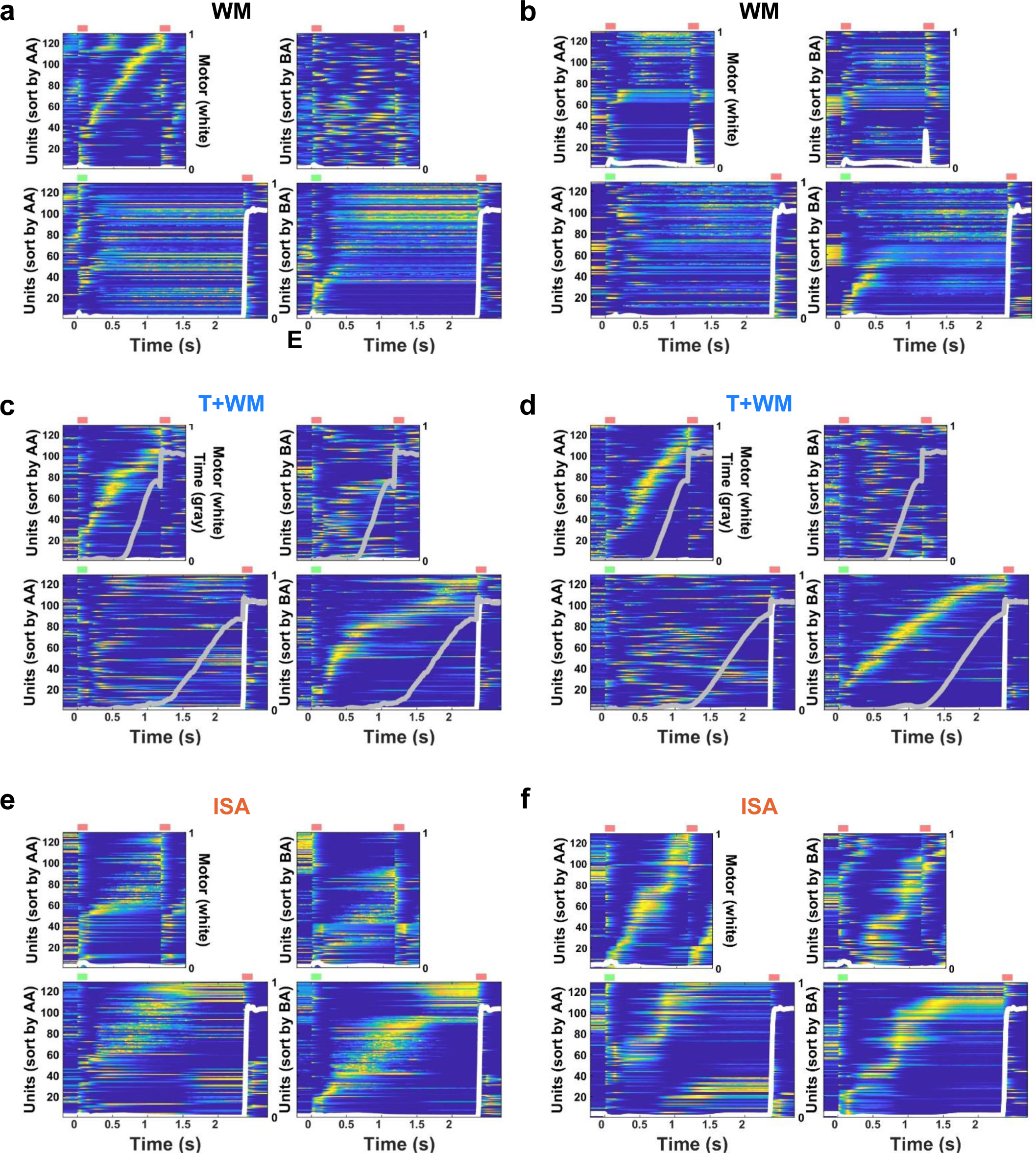
Additional examples of RNN dynamics in the WM, T+WM, and ISA tasks. **a-b,** Two additional sample RNNs trained on the WM task. Panel **a** shows an example of a mixed solution in which cue *A* (short) was encoded in a neural sequence and cue *B* (long) in a fixed-point attractor. **c-d,** Two additional examples of RNNs trained on the T+WM task. **e-f,** Two additional examples of RNNs trained on the ISA task. Note the partial similarity of the patterns evoked by both the *A* and *B* cues. Plots and cross-validated sorting were performed as in Fig. 2. Data correspond to the RNNs included in the analyses of Fig. 2j**-m**.

**Fig. S5.**
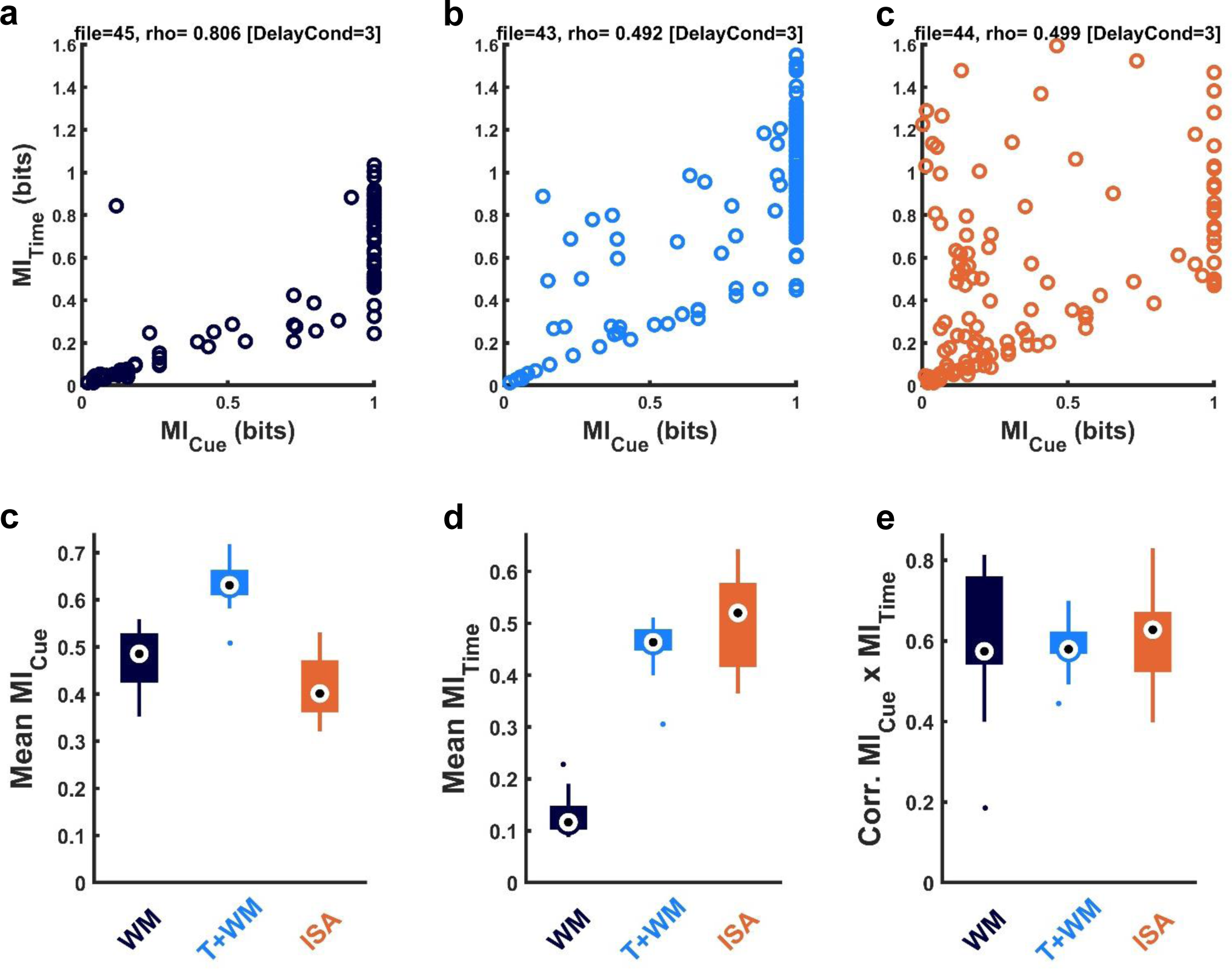
Multiplexing of mutual information for cue and time at the level of individual units. **a,** Sample correlations between the MI for cue and time across units for a sample RNN trained on the WM (left), T+WM (middle), and ISA (right) tasks. The MI values for time were calculated for the long delay, thus only the units that responded more to the long, compared to the short, delay are shown. MI for cue was calculated by averaging the activity over the entire short and long delays. **b,** Medians and interquartile ranges of the mean MI for cue in RNNs trained on each task (n=17 each). **c,** Medians and interquartile ranges of the mean MI for time for RNNs trained on the three tasks. **d,** Medians and interquartile ranges of the Kendall correlation coefficient between the cue and time MI across RNNs. Together plots B-C establish that in the T+WM task individual units contain significant information about Cue (**a**) and elapsed time (**b**), and that there is a strong correlation between them (**c**), indicating multiplexing of WM and time.

**Fig. S6.**
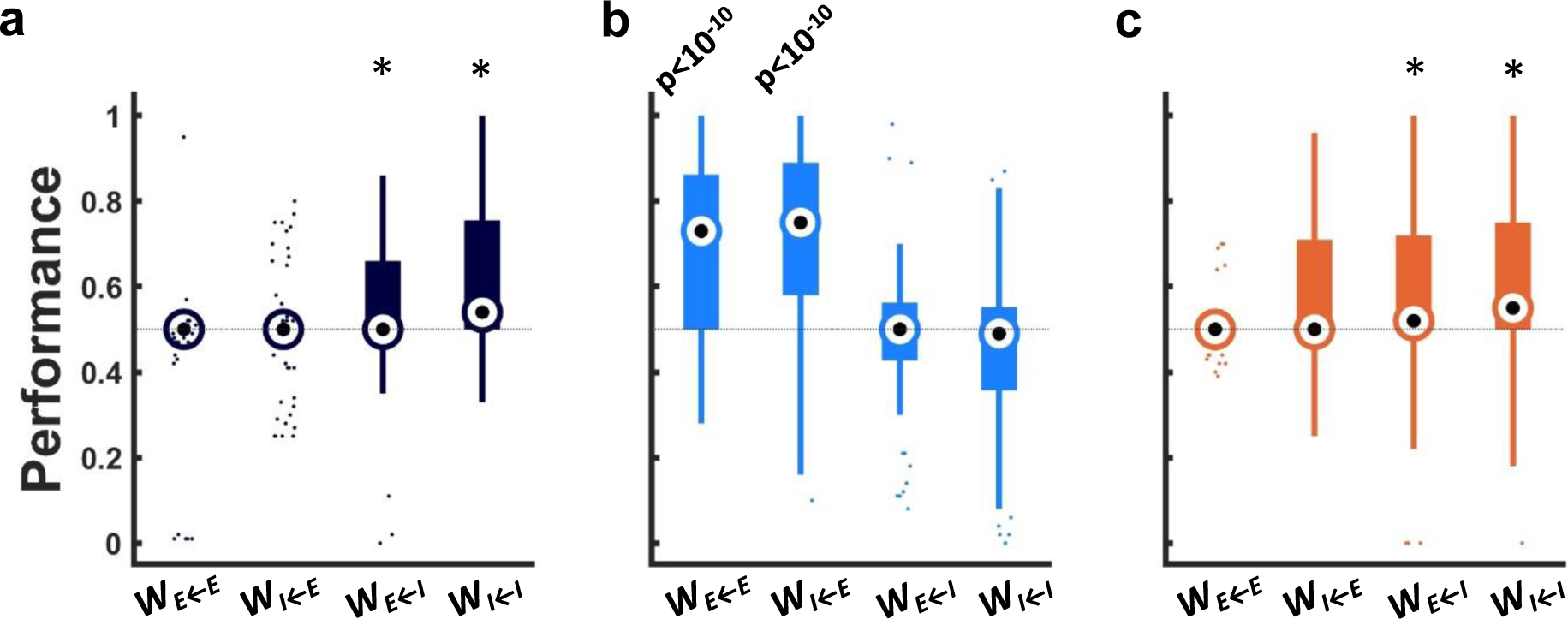
Excitatory connections are the least important in the timing and WM (T+WM) task. Performance after full shuffles of all the nonzero weights of each of the four weight submatrices for the RNNs trained on the WM (**a**), T+WM (**b**), and ISA (**c**) tasks. Performance for the unshuffled RNNs was 100% (Fig. 2j). Shuffling either the W^EE^, W^IE^, W^EI^, or W^II^ matrices resulted in a catastrophic drop in performance to median values close to chance (50%) for both the WM and ISA tasks. Indicating that the synaptic structure of each sub-weight-matrix is critical to the dynamic regimes underlying the computations. In sharp contrast, for the T+WM task shuffling of the excitatory connections (WEE or WIE) did not result in a catastrophic drop in performance, i.e., even after shuffling these weight matrices median performance was approximately 75%, but shuffling inhibitory weights (W^EI^ or W^II^) dropped performance to close to chance. Thus, the excitatory weights were significantly less important than the inhibitory weight for the underlying computations. Asterisks (*) represent p values less than 10^-3^ (Sign rank test against values of 0.5).

**Fig. S7.**
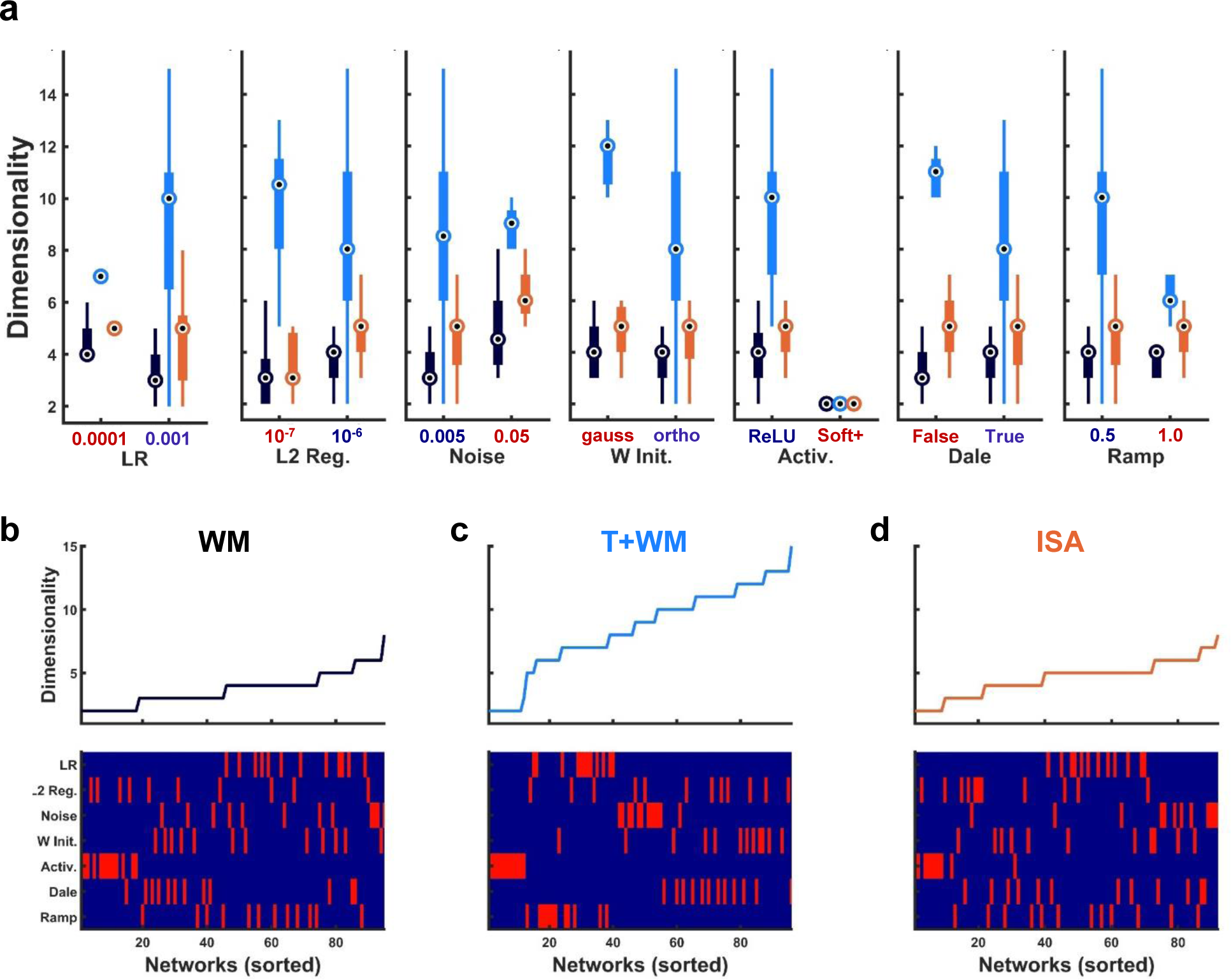
Dimensionality during the delay is higher in the T+WM tasks across a range of hyperparameters. **a,** Dimensionality across seven hyperparameters for the WM, T+WM, and ISA tasks. Blue labels on the x-axis correspond to the default hyperparameters. **b-d,** Relationship between dimensionality in the individual RNNs and the hyperparameters. RNNs were sorted according to their dimensionality across all the hyperparameter runs for the WM (**b**), T+WM (**c**), and ISA (**d**) tasks (upper panels), the corresponding hyperparameters of each RNN in the upper plots are shown in the respective lower panels. Blue values correspond to the default values, and red to the changed hyperparameters as in **a**. Note, for example, that for the T+WM task all the low dimensionality values correspond to the cluster of solid red in the Activation hyperparameter row. Seven hyperparameter values were individually varied in relation to the default hyperparameter regime (8 hyperparameter sets x 3 tasks x 12 seeds = 288 RNNs). Five RNNs did not converge to a performance level > 90% (4 in the ISA task, and 1 in the WM task) and were excluded from this analysis. Note that the ramp hyperparameters do not apply to the WM and ISA tasks (as there is no temporal expectation output unit)—thus the black and orange bars in the ramp panel in **a** corresponds to equivalent simulations.

**Fig. S8.**
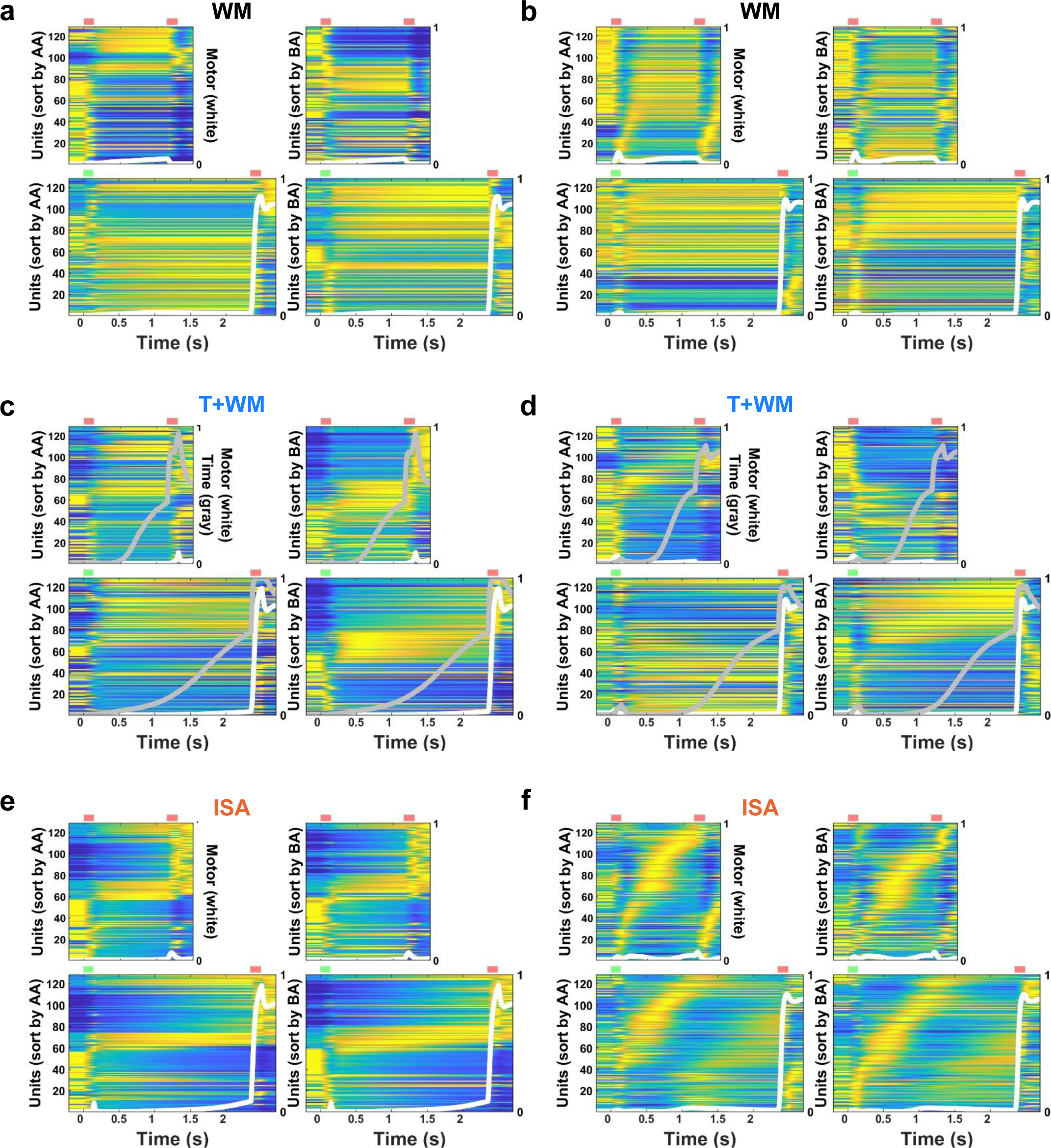
Additional examples of RNN dynamics in the WM, T+WM, and ISA tasks in RNN trained with a softplus activation function. **a-b,** Two RNNs trained on the WM task from the dataset in **Fig S4**. As in Figs. 2 and S4, the diagonal (upper left and lower right) panes are self-sorted (and cross-validated) and the off diagonals are cross-sorted. **c-d,** Two sample RNNs trained on the T+WM task. **e-f,** Two sample RNNs trained on the ISA task. Note that in comparison to the ReLu activation functions maximal activity for many units occurs during baseline, and the presence of both increasing and decreasing ramps.

**Fig. S9.**
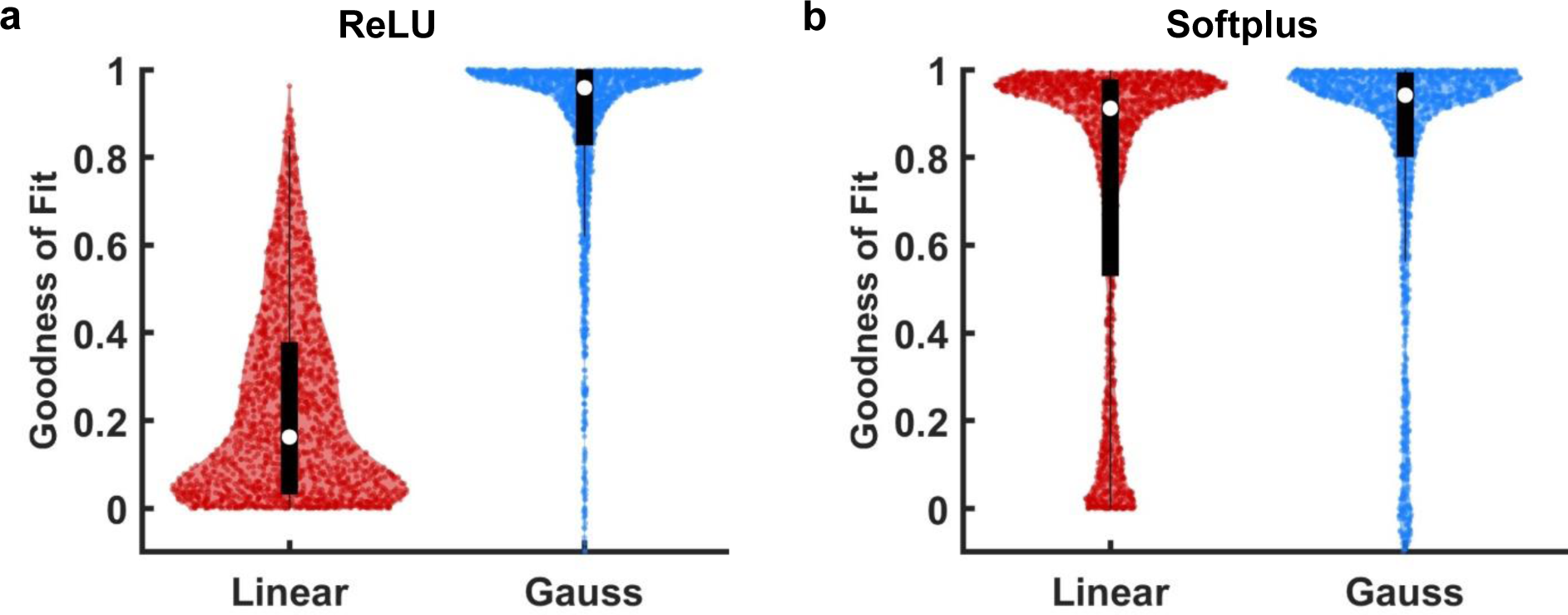
Activation functions (ReLU x Softplus) dramatically alter the encoding of time and WM. **a,** Goodness of fit values for RNNs in the hyperparameter study with ReLU activation functions. The upper half of the most active units during the delay epoch RNNs (n=13) trained on the T+WM task were fit with a linear function (red) or a Gaussian (blue). The degree of freedom was two for both the Linear (slope, intercept) and Gaussian (mean, width) fits. **b,** Corresponding fits for the units in the RNNs with Softplus activation functions. Note that the vast majority of units were well fit with a linear function and that the rising phase of an unconstrained Gaussian function can also fit ramps well. Additionally, note that goodness-of-fit R^2^ values can be negative if a nonlinear fit accounts for less variance than a horizontal line (for visualization purposes only units with R^2^>-0.1 are shown).

**Fig. S10.**
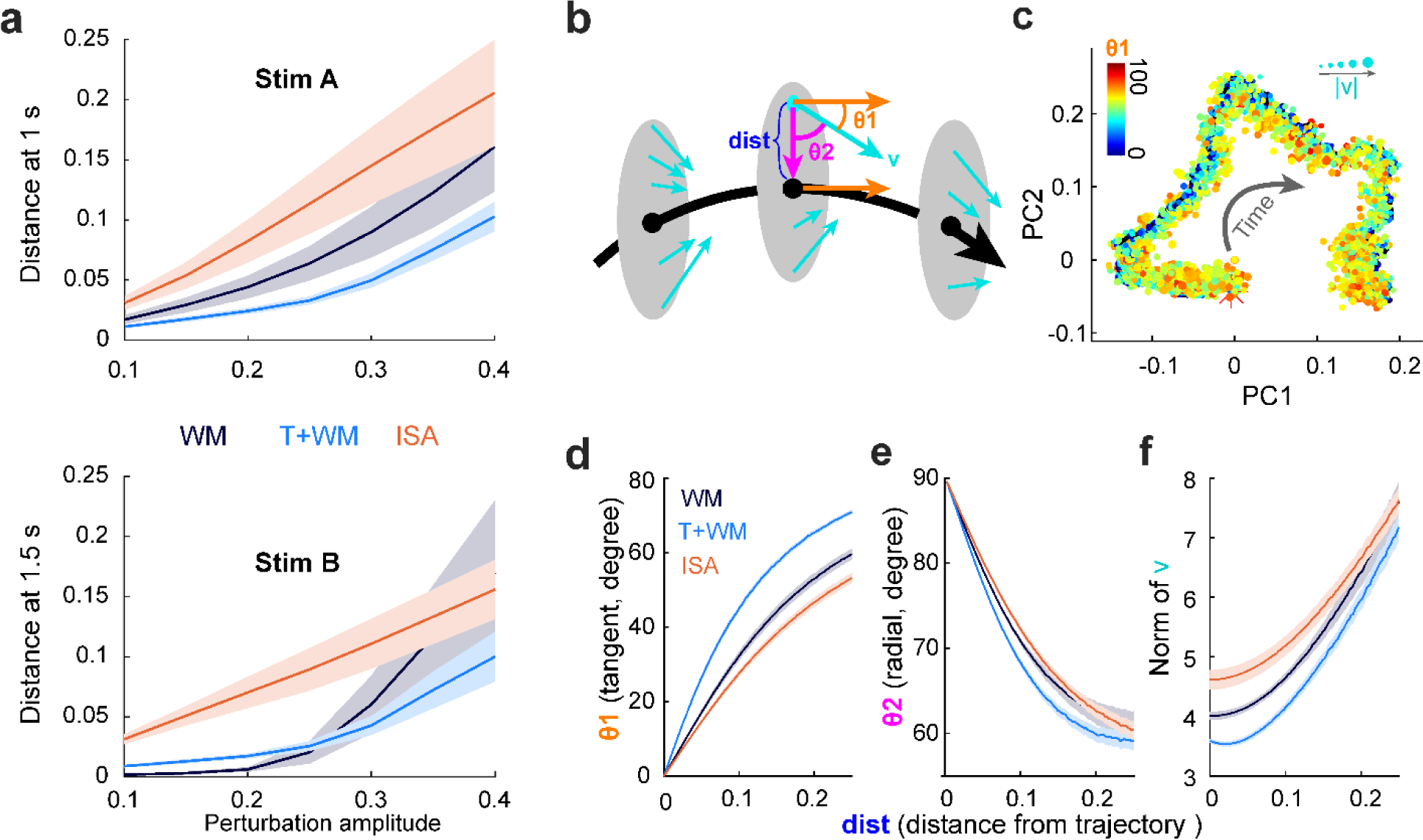
Stability analysis of the network dynamics during the delay period. **a,** Effect of perturbation amplitude applied at 500 ms into the delay period at 1 (Short stimulus, top) or 1.5 s (Long Stimulus, bottom). The Euclidian distance corresponds to the distance between the perturbed and unperturbed trajectories (frozen noise). Note that the perturbation effects are significantly worse in the ISA task, and similar in both the T+WM and WM tasks, even though the WM dynamics approximates a fixed-point attractor. **b**, Schematic of the quantification of the perturbation vector fields. For a given trajectory (black curve) to be a dynamic attractor, the directions of **θ1** (in relation to the tangent direction, orange vector) and **θ2** (the radial direction, magenta vector) are both smaller than 90 degrees. **c**, Visualization of the velocity fields for an example T+WM RNN. At each time point (**r(t)**) on the trajectory (in the full neural space), we sampled 20 points around **r(t)** from a disk (radius of 0.25) orthogonal to the tangent vector of the trajectory. The velocity vectors (**v**) were computed for such points according to the firing rate equation, along with **θ1**, **θ2**, and |**v**|. Finally, the trajectory (black curve) and all the sampling points (round dots) were projected to the two-dimensional PC space. **θ1** and |**v**| for each **v** were then coded by the color and size of the dots respectively. **d**-**f**, Summary for the means of **θ1** (**d**), **θ2** (**e**), and |**v**| (**f**) at time 1.5 s (1 s after the perturbation) vs. the amplitude of the perturbation (distance to the reference trajectory) from 17 RNNs in the WM, T+WM, and ISA tasks.

